# Three Novel Methods for Determining Motor Threshold with Transcranial Magnetic Stimulation Outperform Conventional Procedures

**DOI:** 10.1101/2022.06.26.495134

**Authors:** Boshuo Wang, Angel V. Peterchev, Stefan M. Goetz

## Abstract

**Objective:** Thresholding of neural responses is central to many applications of transcranial magnetic stimulation (TMS), but the stochastic aspect of neuronal activity and motor evoked potentials (MEPs) challenges thresholding methods. We analyzed existing methods for obtaining TMS motor threshold and their variations, introduced new methods from other fields, and compared their accuracy and speed.

**Approach:** In addition to existing relative-frequency methods, such as the five-out-of-ten method, we examined adaptive methods based on a probabilistic motor threshold model using maximum-likelihood (ML) or maximum *a-posteriori* (MAP) estimation. To improve the performance of these adaptive estimation methods, we explored variations in the estimation procedure and inclusion of population-level prior information. We adapted a Bayesian estimation method which iteratively incorporated information of the TMS responses into the probability density function. A family of non-parametric stochastic root-finding methods with different convergence criteria and stepping rules were explored as well. The performance of the thresholding methods was evaluated with an independent stochastic MEP model.

**Main Results:** The conventional relative-frequency methods required a large number of stimuli, were inherently biased on the population level, and had wide error distributions for individual subjects. The parametric estimation methods obtained the thresholds much faster and their accuracy depended on the estimation method, with performance significantly improved when population-level prior information was included. Stochastic root-finding methods were comparable to adaptive estimation methods but were much simpler to implement and did not rely on a potentially inaccurate underlying estimation model.

**Significance:** Two-parameter MAP estimation, Bayesian estimation, and stochastic root-finding methods have better error convergence compared to conventional single-parameter ML estimation, and they all require significantly fewer TMS pulses for accurate estimation than conventional relative-frequency methods. Stochastic root finding appears particularly attractive due to the low computational requirements, simplicity of the algorithmic implementation, and independence from potential model flaws in the parametric estimators.

## 1. Introduction

### 1.1. Significance and challenges of thresholding in TMS

Transcranial magnetic stimulation (TMS) is a noninvasive technique to activate neurons in the brain [1], [2]. Certain pulse rhythms can further modulate neural circuits, i.e., change how they process endogenous signals [3]. A key parameter of the stimulation pulses is their strength, which influences not only how many neurons but also which neuron types are stimulated [4]–[6]. As the TMS pulse strength affects the activation of neuron types with different functions, it also influences the efficacy of neuromodulation protocols and can even reverse the direction of the effect [7]–[11]. Finally, the stimulation strength is a key parameter not only for ensuring an effective and reproducible procedure but also for safety, and strength limitations for various modulatory protocols are recommended to avoid seizures [12]–[14].

Typically, the TMS pulse strength is individualized to a specific subject’s or patient’s anatomy and physiology based on a measured response threshold. A commonly used threshold is the motor threshold based on activation of finger muscles to stimulation of their representation in the primary motor cortex, but other types of thresholds could also be used, such as phosphene threshold of visual artifacts elicited in the visual cortex, threshold of cortical silent period, and transcranial evoked potentials recorded from electroencephalography (TMS-EEG) [15]–[22]. For the motor threshold, the muscle movement following stimulation in the primary motor cortex is observed visually in some cases, typically in clinical practice, whereas in other cases, typically in research, the size of motor evoked potentials (MEPs) is measured with electromyography (EMG) at the target muscle [23]–[25]. The motor threshold is routinely also used as a reference for other brain targets and its detection is recommended before any further neuromodulatory intervention, even for recurring subjects or patients [26]–[28]. Furthermore, the threshold and its changes can serve as a quantitative metric, e.g., for the diagnosis of lesions [29]. The centrality of the motor threshold to TMS procedures underscores the need for a well-defined, rapid, accurate, and unbiased thresholding procedure.

However, the MEPs in response to stimulation in the primary motor cortex are highly stochastic due to the dynamic variability in cortical neurons and the corticospinal tract—a pulse administered twice with identical strength can lead to a maximum MEP response close to saturation once and no response at all in the other case [30]– [32]. Moreover, the variability is far from GAUSSIAN and appears to involve several variability sources [32]–[35]. Consequently, accuracy of the estimation procedure is hard to guarantee and achieving it with a suboptimal estimation algorithm can be time consuming.

### 1.2. Existing motor threshold estimation methods

The TMS motor threshold, *x*_th_, is usually defined as the lowest stimulation pulse amplitude, *x*, given as a percentage of the maximum stimulator output (MSO), leading to a peripheral muscle response, *r*, with a 50% probability [27], [28], [36],

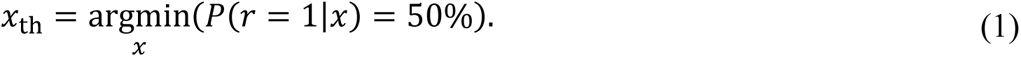

The binary response *r* is defined as

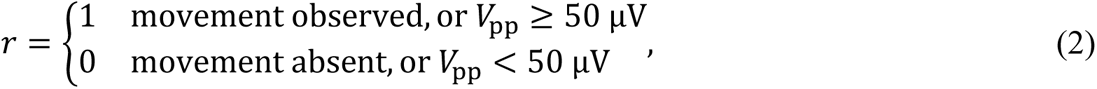

where *V*_pp_ is the peak-to-peak EMG amplitude [37]. Historically, the value of 50 μV was chosen relatively arbitrarily as a low level that is not far from the noise limit of most EMG detection systems [38]–[40]. The most common practice for motor threshold finding, or “threshold hunting”, is estimating the relative frequency of positive and negative responses by appropriate stepping of the stimulus amplitude to find the 50% probability level. Therefore, at each test pulse amplitude, a number of stimuli—usually up to ten—have to be administered, hence the name “five-out-of-ten method”. Such an approximation of the probability by a relative frequency within just ten stimuli can be rather imprecise, and a larger number of trials, e.g., twenty or more, per stimulation amplitude is required to obtain more reproducible results for clinical and research purposes [28], [41]. Given its simplicity, the relative-frequency method is widely used and often serves as a benchmark for other methods, despite concerns regarding its mathematical validity or usefulness [41]–[43].

Another well-described method based on the relative response frequency was proposed by MILLS and NITHI [39]. Instead of detecting the threshold directly, it uses one point above threshold (*upper threshold*) and one below (*lower threshold*), and the threshold estimate itself is defined as the mean of these two points. Mathematically, this relates to a local linearization of the sigmoid curve of the neuronal response in dependence of the stimulation amplitude in the probability space. Like the relative-frequency method, both upper and lower thresholds are defined statistically—all ten responses suprathreshold and subthreshold, respectively. Both the relative-frequency and MILLS-NITHI methods require a high number of stimuli [40], [44] and consequently a long duration to complete due to the necessary pause between two stimuli for preventing inter-pulse correlation [45]–[47]. Furthermore, a large number of stimuli also bears a risk of unintended neuromodulation and side effects [14].

Instead of determining relative frequencies, an alternative thresholding method performs a maximum-likelihood (ML) test for a predefined model describing the sigmoidal probability distribution of the binary response as a function of stimulation amplitude—most commonly of a cumulative GAUSSIAN function with two parameters, the midpoint/threshold and slope/spread^1^. This maximum likelihood estimation (MLE) method was introduced by Awiszus [48] and was adapted from psychophysics where responses have strong stochastic characteristics as well [49], [50]. The result of such parameter estimation by sequential testing (PEST) provides the most likely parameters, which include the threshold. An essential condition is a good selection of the tested stimulus amplitudes [51]. In contrast to frequency-based methods, each amplitude is usually applied just once. Mishory et al. [44] reported that the number of required stimuli and time for determining threshold was markedly lower for ML-PEST compared to the MILLS–NITHI two-threshold method, whereas both methods have similar accuracy.

### 1.3. Limitations of existing methods and alternative approaches

Existing motor threshold estimation methods have some limitations that can be addressed to improve the accuracy and speed of the procedure. A key feature that can notably influence performance is the stepping rule [51], which chooses the next stimulation amplitude adaptively by taking into account the information from the previous observations. For the next stimulation amplitude, the ML-PEST [48] chooses the MLE of the threshold, which is obtained from a reduced single-parameter model of the response and uses a fixed nonstochastic relationship for the spread of the probability function. This relationship is assigned based on averages from representative measurements [48], and therefore does not reflect individual variability. A limitation of this approach is that by not allowing any deviation from a fixed spread–threshold relationship, the method may place all stimuli systematically too high or too low with respect to the optimal stepping for an individual subject, which can lead to slow stepping and large threshold misestimations if the initial stimulation amplitude is poorly chosen [52], [53].

The threshold estimation can be improved by a full two-parameter MLE in which both the threshold and spread are free to vary. Incorporating additional statistical information from prior or typical results of the thresholding procedure can further improve the estimation [52], [54], which is a method adapted from the field of psychophysics [51]. Although Qi et al. called it a BAYESIAN method [54], this name is not adopted here since the approach incorporates *a-posteriori* information into a classical FISHER MLE using BAYES’ theorem. This has been known in mathematics [55] and applied fields [56] as maximum *a-posteriori* (MAP), a term we adopt in this paper.

While BAYES’ theorem has been used for MAP estimation to improve the TMS threshold estimate [52], [54], both ML and MAP methods are point estimators. In contrast to these point estimators, strictly BAYESIAN methods that are consequently and completely premised on BAYESIAN statistics [57]–[59] have not been used in TMS. In this text, the BAYESIAN style particularly refers to the updating rule for the probability function after new observations, which should therefore follow the BAYESIAN learning paradigm. We adapted the BAYESIAN estimation method from KONTSEVICH and TYLER [57] for TMS motor threshold estimation, using the same model as in the point estimators.

Both the relative-frequency methods and the parameter estimation methods adapted from psychophysics reduce the TMS motor responses to binary events (2). Whereas both psychophysics and TMS motor threshold determination by visual observation of finger twitches deal only with binary responses to a stimulus (perceived or not perceived), the use of EMG for measuring muscle activation by TMS provides a continuous response strength and thus richer information. The EMG motor threshold is not a binary transition between no response and response but a point on the dose–response curve close to the lowest stimulation strength that elicits an excitatory response detectable with conventional EMG recording and analysis methods [60]–[62]. Further, the motor threshold is likely not the lowest stimulation strength to evoke detectable responses, as recent studies demonstrate that the 50 µV threshold is likely closer to the middle of the logarithmic dose–response curve of MEPs when it is separated from the baseline noise [63], [64]. The smoother transition of the expected value of the responses in dependence of the stimulation strength may allow the use of the MEP amplitude value to find the threshold faster. To consider this additional analog information of the response amplitude appropriately, stochastic root-finding methods [65], [66] (also known as stochastic approximation) are adapted and evaluated here.

### 1.4. Organization of this paper

We first describe existing TMS thresholding methods as well as the aforementioned improvements and introduce the BAYESIAN estimation and stochastics root-find methods, covering a total of three classes and 25 variations of methods. These methods are divided into two categories, based on whether a parametric model describing the motor response underlies the method. To evaluate and compare the thresholding methods, an independent testbed incorporating a detailed stochastic MEP model [61] is used, with 25,000 virtual subjects. The performance of the thresholding methods is analyzed based on their accuracy of the threshold estimate and speed to acquire threshold (relative-frequency methods) or convergence rate of the threshold error (other methods). Both the relative and absolute errors serve for a comparison of the accuracy, and statistics including the median, the upper and lower quartiles and adjacent values, and outliers were calculated for the virtual population. The best performing method of each class are identified and the trade-off between different methods is discussed. The results show that the three alternative approaches—two-parameter MAP estimation, Bayesian estimation, and stochastic root-finding—outperform single-parameter ML estimator, and they all require significantly fewer TMS pulses for accurate motor threshold estimation than the conventional relative-frequency methods.

## 2. Methods

### 2.1. Overview of thresholding methods

The estimation of a neuronal property such as the excitation threshold can be performed in two fundamentally different ways. The model-based parameter estimators (ML, MAP, and the BAYESIAN methods) require a probabilistic description of the input–output black box being analyzed in dependence of a certain parameter set. An inaccurate model will very likely deteriorate the quality of the estimate. In contrast, the early approaches, namely the five-out-of-ten or ten-out-of-twenty relative-frequency method based on recommendations of the International Federation of Clinical Neurophysiology (IFCN) [27], [28] or the MILLS–NITHI two-threshold method [39], do not need such a predefined model description as they refer directly to the definition of the threshold. The stochastic root-finding methods can also be counted among this class of non-parametric systems. In the following, the existing and several new or adapted algorithms are shortly described and compared for the application in TMS. Computation and data analysis were performed with MATLAB (versions 2021a and 2023a, The Mathworks, Natick, MA, USA) on the Duke Compute Cluster, with the code of the thresholding methods available at [67].

### 2.2. Nonparametric methods

#### 2.2.1. IFCN relative-frequency method

The procedure described by the IFCN [27], [28], [68] leaves freedom of interpretation of some details. For the analysis in this paper, the relative-frequency method was defined as follows to realistically represent the implementation in many laboratories:

a) The starting amplitude is known from the hotspot search (see section 2.4.2 Simulation routine for further details on the starting amplitude), in which the corresponding neuron population being responsible for activating the observed muscle group (e.g., the *first dorsal interosseus* or the *abductor pollicis brevis*) is located with test stimuli on the primary motor cortex. The required amplitude adjustments in this step lead to an amplitude not too far above the desired threshold as spatial focality would be compromised with stronger stimuli.
b) At a given stimulation amplitude, a number of stimuli are applied up to a maximum of ten or twenty until either the count of positive responses reaches half the number (five or ten, respectively) or that of negative responses exceed half the number (six or eleven, respectively).
c) Being suprathreshold by definition in the situation of positive responses accounting for at least half the number of stimuli, the stimulation amplitude is stepped downward, typically by 1% or 2% MSO [28], [68], and Step b) is repeated.
d) The latter outcome in Step b) precludes 50% positive responses within the number of stimuli and the amplitude is subthreshold. The threshold itself is defined as the last level before this termination point.

Versions with the number of stimuli of ten and twenty and step size of 1% and 2% MSO were performed to show the effect of the number of stimuli per step and the step size on the accuracy and the number of stimuli required for the methods (Table 1).

**Table 1.** Summary of relative-frequency-based methods

| Method | Variation | Step size (% MSO) |
| --- | --- | --- |
| IFCN relative-frequency | Five-out-of-ten | 1 |
|  |  | 2 |
|  | Ten-out-of-twenty | 1 |
|  |  | 2 |
| MILLS–NITHI | Original | 1 |
|  |  | 2 |
|  | Modified | 1 |
|  |  | 2 |

#### 2.2.2. MILLS–NITHI two-threshold method

The original MILLS–NITHI method [39] was carried out as follows:

a) The subject is stimulated with pulses starting at an amplitude of 20% MSO and increasing in 10% MSO steps until the first positive response is identified.
b) Applying up to 10 pulses per stimulation amplitude, the stimulation is then decreased in 1% MSO steps when a positive response is observed. The intensity at which all 10 stimuli produce no response is designated as the lower threshold.
c) Starting from the lowest amplitude that has not resulted in a “no response”, i.e., the last amplitude that has one stimulus with positive response during Step b), and increasing the stimulation in 1% MSO steps, the upper threshold is designated as the minimum intensity at which 10 stimuli all produce positive responses.
d) The threshold is calculated as the mean of the lower and upper thresholds.

In our study, a modified version of the MILLS–NITHI method was performed using the same starting amplitude as the IFCN relative-frequency method. The first negative response was identified by decreasing the amplitude in 10% steps and the LT and UT were then similarly obtained as in the original methods but in reversed order. Step sizes of 1% and 2% MSO were tested (Table 1).

#### 2.2.3. Stochastic root-finding methods

The IFCN relative-frequency and MILLS–NITHI methods described above both require binary information reduction through quantization of the response (above versus below 50 μV). In contrast to such quantization, robust numeric root-finding algorithms that are still relatively unknown in TMS can directly act on analog data, e.g., the voltage amplitude of the EMG response. Preserving valuable continuous information in the response, such root-finding approaches could enable an improvement of convergence. The sequential formulation proposed here was derived from the method of ROBBINS and MONRO [65], [66], with the neuronal variability playing the role of a noise source. The requirements for this approach are rather weak and comprise constant monotonicity on average [65].

The next stimulation amplitude, *x*_*i*+1_, was defined relative to the current one using an iterative analog control sequence (ACS) that was intended to converge towards the threshold:

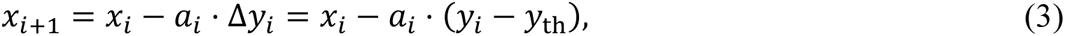

where *i* ≥ 1 was the step number, *a*_*i*_ was the step size of the control sequence, *x*_*i*_ denoted the stimulation amplitude, and *y*_*i*_ and *y*_th_ were logarithms of the response (*y*_*i*_ = lg(*V*_*i*_/[1 V])) observed at step *i* and at the real threshold *V*_th_ (50 μV peak-to-peak by definition), respectively. The log-transformation accounted for the stochastic characteristics of EMG responses and the first-approximation log-normal distribution of the response amplitude following an inductive stimulus in the motor cortex [46]. The resulting distribution of *y*_*i*_ had reduced skewness and should be closer to GAUSSIAN and therefore statistically well-behaved. Performing the whole procedure in the logarithmic space also granted the requirement of constant monotonicity. The iteration for the EMG response thus became

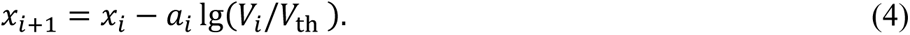

We implemented several versions of the sequence of step sizes *a*_*i*_ to compare the convergence properties (Table 2). In the basic version ACS1-H, the step size followed a harmonic series with almost sure (a.s.) convergence^2^

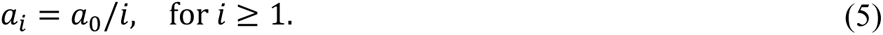

**Table 2.** Summary of root-finding methods

| Method | Control sequence | Order | Convergence of $a_i$ |
| --- | --- | --- | --- |
| ACS1-H | Analog | 1 | Harmonic ($i^{-1}$), non-adaptive |
| DCS1-H | Digital | 1 |  |
| ACS2-H | Analog | 2 |  |
| ACS1-HA | Analog | 1 | Harmonic ($i^{-1}$), adaptive |
| DCS1-HA | Digital | 1 |  |
| ACS2-HA | Analog | 2 |  |
| ACS1-GA | Analog | 1 | Geometric ($2^{-i/2}$), adaptive, |
| DCS1-GA | Digital | 1 |  |
| ACS2-GA | Analog | 2 |  |
| ACS1-GHA | Analog | 1 | Geometric ($2^{-i/2}$) switching<sup>a</sup> to harmonic ($i^{-1}$), adaptive |

<sup>a</sup>Transition when the maximum change of $x_i$ , i.e., $|\Delta x_i| = |a_i \cdot \Delta y_i|$ , is expected to be less than 1.5% MSO.

Following the style of Delyon and Juditsky [69], adaptive methods were utilized to accelerate convergence by adjusting the control sequence at step *i* (*i* > 1) only when the response changed from subthreshold to suprathreshold or vice versa from the previous step (*i* − 1), i.e., following a sign change in Δ*y*_*i*_: sgn(Δ*y*_*i*_) ≠ sgn(Δ*y*_*i*−1_). For harmonic convergence, the corresponding adaptive control sequence ACS1-HA was

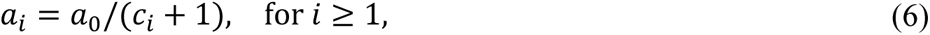

where *c*_*i*_ was the number of sign changes of Δ*y*_*i*_ up to step *i*,

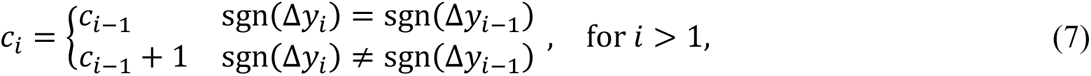

with *c*_1_ ≜ 0. Unlike classical mathematical problems, the TMS sequence has only a very limited number of steps. Thus, quick convergence seems more attractive here rather than any statement for the case of infinite duration. To test faster geometric convergence of 2^−*i*/2^, ACS1-GA used an adaptive control sequence of

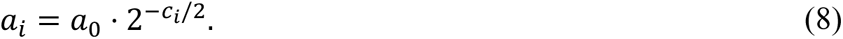

Further variations of these methods were designed to examine the convergence rate and performance. To take advantage of both fast convergence and the a.s. convergence properties, a hybrid method ACS1-GHA was designed to switch the control sequence *a*_*i*_ from geometric to harmonic convergence when the threshold estimate was close to the true threshold and the expected change in *x*_*i*_ (|Δ*x*_*i*_| = |*a*_*i*_ ⋅ Δ*y*_*i*_|) was small—less than 1.5% MSO. Quantized versions, i.e., digital control sequences (DCS), were implemented for comparison as well. The analog value Δ*y*_*i*_ was reduced using a modified sign function to δ*y*_*i*_ = sgn(Δ*y*_*i*_) ∈ {−1, +1} for the two different cases of below threshold and above (or equal) threshold. DCS allowed the stochastic root-finding method to be used also without EMG recordings, just “response” and “no response” inputs, in a manner compatible with the other estimation methods. Second-order versions of the analog control sequences (ACS2) were also implemented, in which the next stimulation amplitude was defined using information from the previous two steps

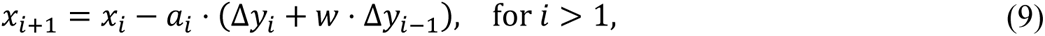

with *w* being the weight for the second-order term.

The definition of appropriate initial step size *a*_0_ was assisted by the iteration formulation (3), which resembled a slope [70] after rearrangement

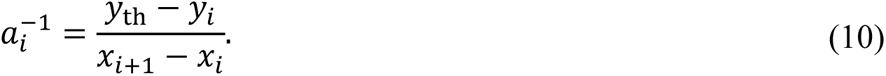

For reaching the threshold quickly, *a*_0_ should accordingly be not too far from the cotangent of the line connecting the present stimulation–response (*x*_*i*_, *y*_*i*_) towards the threshold (*x*_th_, *y*_th_). Larger values increase the speed of approach of *x*_*i*_ towards *x*_th_, but may cause the stepping to overshoot and oscillate around the target, whereas smaller numbers promote lengthy one-sided convergence. Statistical information about typical and maximum slopes can inform the choice of *a*_0_. For the simulations in this paper, the default initial step size was chosen such that it approximated typical cotangents around the inflection point of the recruitment curve (see section 2.4.1 Response model)

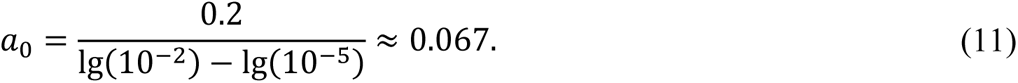

Here lg(10^−5^) and lg(10^−2^) (i.e., 10 μV and 10 mV, respectively) were approximate response amplitudes near the EMG noise floor and saturation level at the lower and higher ends of the input–output curve, respectively, and 0.2 (20% MSO) was a sufficient change in stimulus strength to cover the transition of the typical sigmoidal response curve. Initial step size from one tenth to ten times the default values evenly sampled on a logarithmic scale were also tested. In second-order sequences, the weight of the second-order term also affects convergence. Larger/positive *w* speeds up the approach to threshold, but could generate oscillations depending on the value of *a*_0_; smaller/negative *w* reduces the speed of convergence, but could help to mitigate the oscillation if *a*_0_ is chosen too large. A default value of −0.1 was used and values between −1 and 1 were tested for ACS2-HA.

### 2.3. Parametric methods

#### 2.3.1. Basic maximum-likelihood estimation method

The implementation of the basic MLE method (ML-PEST) followed substantially the suggestions and definitions by Awiszus [48]. The probability space was described by a cumulative GAUSSIAN model with two degrees of freedom—the threshold *t* = *x*_th_ and the spread *s*:

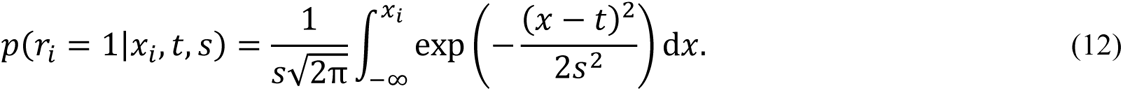

Here, *r*_*i*_ denoted the binary response (2) of a single stimulus *x*_*i*_, and the slope at the threshold, 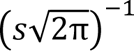, was inversely related to the spread.

The likelihood function was defined for independent events

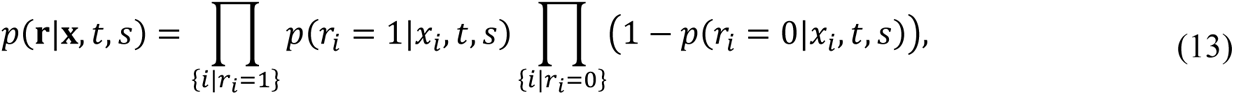

where **x** and **r** were the lists of stimuli and corresponding responses, respectively. According to the classical FISHER approach, the maximum of this expression for the arguments *t* and *s* provided the most likely parameter set (argmax_(*t*,*s*)_ *p*(**r**|**x**, *t*, *s*)). Due to the strict monotonicity of the logarithm, the product can be split into an equivalent sum, which formed the log-likelihood and was also numerically more robust. The MLE of *t* provided both the threshold estimate at the current step and stimulation amplitude at the next step.

Previous implementations of MLE (13) (e.g., MTAT and ATH-tool, [48], [71], [72]) were simplified by using only the threshold parameter *t*, whereas the spread parameter was fixed to the value *s* = 0.07 ⋅ *t*. We replicated this method (MLE-1) and also performed a full MLE with both parameters (MLE-2) (Table 3). Two approaches were implemented and compared to find the threshold *t* that maximize the likelihood function. The first approach directly maximized the likelihood function over *t* (using MATLAB function `fmincon`, applied on the negative of the log-likelihood). This approach, however, could be unstable during the initial steps when there were few amplitude and response samples (**r**, **x**) and the likelihood function had a plateau, i.e., a range of *t* or (*t*, *s*) combinations resulted in the same maximum. The second approach explicitly evaluated the likelihood function and calculated the mean of the argument *t* over the range of the plateau [48]. For the explicit evaluation, the parameter space was discretized with Δ*t* = 0.002 for *t* ∈ (0, 1.3], and 100 logarithmical-distributed samples *s* were chosen from [0.005, 0.5] for MLE-2. The upper bound of the threshold parameter exceeded one, which is explained in section 2.4.1 Response model.

**Table 3.** Summary of parametric estimation methods

| Method | Threshold estimation | Stepping |
| --- | --- | --- |
| MLE-1<sup>a</sup> | Single-parameter MLE |  |
| MAP-1 | Single-parameter MAP |  |
| MLE-2<sup>a</sup> | Two-parameter MLE |  |
| MAP-2 | Two-parameter MAP |  |
| BAYESIAN, BAYESIAN-NP<sup>b</sup> | Expectation of threshold | Minimum expected entropy |
| BAYESIAN-TS | Expectation of threshold |  |

<sup>a</sup> Implemented using both direct maximization and explicit evaluation of the log-likelihood function.
<sup>b</sup> Implemented with no prior population information.

#### 2.3.2. Maximum-a-posterior estimation method

Modified estimation methods (MAP-1 and MAP-2, Table 3) were designed based on a MAP-like approach which inversed the likelihood ratio in (13) [52], [54] and directed the likelihood function backwards from the perspective of the causality of the model. For incorporating statistical information, BAYES’ theorem was used to modify the likelihood function (13) to improve the performance for common cases per

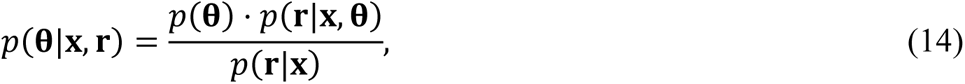

in which the vector **θ** = (*t*, *s*) denoted the parameter set. The additional information about the distribution of typical parameters *p*(**θ**) had to be provided in advance (see section 2.4.1 Response model). Despite that, this was not *a-priori* information from an epistemological point of view, since the algorithm can be designed to collect the information itself. The last factor *p*(**r**|**x**) can be evaluated using

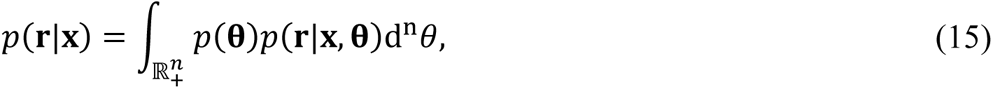

with *n*=2 being the number of parameters. For maximizing the logarithm of (14) with respect to **θ**, however, the denominator *p*(**r**|**x**) was a constant and therefore was omitted.

#### 2.3.3. Kontsevich–Tyler Bayesian estimation method

The BAYESIAN methods (Table 3) here were adapted from KONTSEVICH and TYLER [57]. All central elements were taken as described, using the GAUSSIAN black box (12) as in the other ML and MAP estimators for consistency. Based on this, new information was embedded into the probability density function *p*(*r*|*x*, **θ**) at every step *i* using BAYES’ theorem following

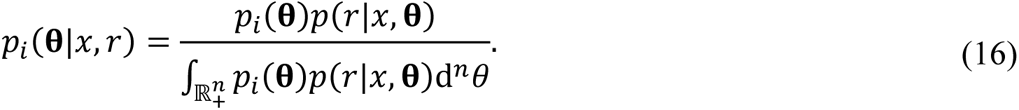

The next stimulation amplitude *x*_*i*+1_ was determined using an information-theoretical approach which minimized the expected entropy at the next step [57]

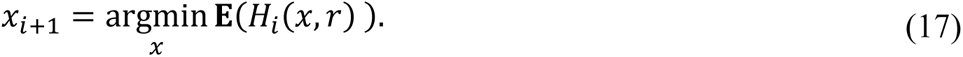

Here, **E** denotes expectation and the entropy *H*_*i*_(*x*, *r*) was defined as

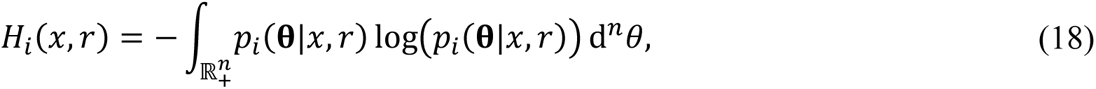

and the expectation was a sum of the entropy *H*_*i*_(*x*, *r*) weighted by the probability *p*_*i*_(*r*|*x*) of sub- and supra-threshold responses

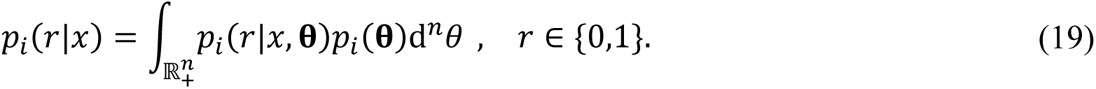

Each stimulus–response pair narrowed down the posterior probability density distribution to

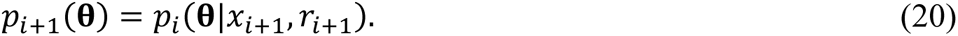

The expected value provided the estimates for the parameters at each step per

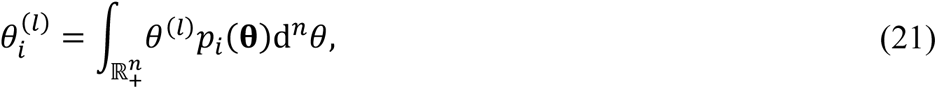

where *l* was the parameter index (i.e., *l* = 1,2 for **θ** = (*t*, *s*)). In contrast to the point estimator methods, the probably density functions (12) and (16) were explicitly computed and stored for the whole parameter space: *r* ∈ {0,1}, *x* ∈ (0,1.3], *t* ∈ (0,1.3], and *s* ∈ [0.005, 0.5] . For computational purposes, the parameter space was discretized with Δ*x* = 0.01 and Δ*t* = 0.002, and 100 logarithmical-distributed samples were chosen for *s*, corresponding to 8.5 million points for *p*(*r* = 1|*x*, **θ**). The initial distribution *p*_0_(**θ**) was chosen as the population distribution *p*(**θ**) (see section 2.4.1 Response model).

Two variations of the BAYESIAN method were implemented. The first one had no prior (NP) population information—using a uniform distribution for *p*_0_(**θ**)—to evaluate the contribution of the prior information to convergence. To compare the entropy minimization stepping rule against the threshold stepping (TS) of the MLE and MAP methods, the second variation skipped the entropy calculation ((17)‒(19)) and used the latter’s choice of the threshold estimate at step *i* (21) for the stimulation amplitude for the next step: 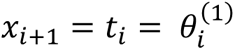

### 2.4. Testbed

#### 2.4.1. Response model

We implemented a realistic MEP model [61] which included variability patterns such as amplifier noise, saturation, and neural variability behavior [30], [34], [35]. The model’s increased complexity compared to other numerical studies concerning threshold-estimation methods [48], [73] was intended not only to reproduce more faithfully MEP behavior but also to avoid using the very same models underlying the parameter estimation methods for the test procedure, which would risk numerical favoritism for those particular methods. The response model provided analog EMG amplitude values based on a stochastic recruitment curve (Figure 1Ai), with parameters mimicking real measurements [74]. The analog amplitude response, i.e., the peak-to-peak MEP amplitude *V*_PP_ in units of volt, was defined as

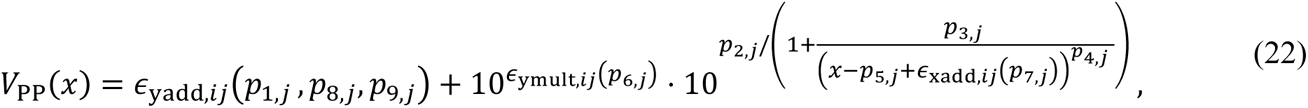

where *p*_1,*j*_ to *p*_9,*j*_ were the individual response parameters for subject *j* obtained from an experimental population [34], and intra-individual variability sources *ϵ*_xadd,*ij*_ and *ϵ*_ymult,*ij*_ and the additive noise *ϵ*_yadd,*ij*_ varied between subjects *j* and for pulses *i* along distributions with subject-dependent parameters. The distributions of recruitment parameters, intra-individual variability, and noise are given in the Appendix. While nominally the stimulation amplitude *x* should be in the range [0,1], the response model was fitted to experimental data obtained with a controllable pulse parameter TMS (cTMS) device using 60 μs pulse width with the maximum capacitor voltage chosen to match stimulation strength to conventional TMS devices for a specific muscle target [34], [74], [75]. Therefore, stimulation amplitudes higher than the nominal 100% MSO could be achieved by increasing the capacitor voltage or with a different TMS device. To focus on the performance of the thresholding algorithms themselves and eliminate influence from the device and choice of stimulation target, we extended the range of possible stimulation amplitudes up to 130% MSO in the model, which was 30% above the maximum threshold within the virtual population.

Model parameters were generated for a total of 25,000 virtual subjects. Each subject’s input–output curve was characterized using 100,000 stimulation–response pairs, with a higher number of pulses applied near the amplitudes that generated a 50 μV response in the corresponding *noiseless* model (see Appendix). The ground truth threshold for each subject was determined by linear regression of the relative frequency of suprathreshold responses to find the amplitude with 50% response probability (Figure 1, panels Aii, Bi, and Bii). The regression also provided the slope at the threshold, which was converted to the spread for the corresponding GAUSSIAN model (Figure 1, panels Bi and Biii). The ratio between the spread and threshold (Figure 1Biv, median, mean, and SD of 0.076, 0.077, and 0.012, respectively) was close to but on average ∼ 10% larger than the fixed nominal value of 0.07 from a previous study [48] and also had a wide distribution. The probability density distribution of threshold and spread *p*(*t*, *s*) can be fitted with normal and lognormal distributions, respectively, with details given in the Appendix. This distribution *p*(*t*, *s*) served as prior information *p*(**θ**) or *p*_0_(**θ**) for the MAP and BAYESIAN methods, respectively. By applying MLE to each subject, the parameters *t* and *s* for the Gaussian model were obtained and analyzed (Figure S1A). For approximately 90% of subjects, the response probability distributions estimated with GAUSSIAN MLE matched their ground truth well (Figure S1B). For the remaining 10% of subjects, however, MLE resulted in large errors (especially for the spread parameter), showing that the GAUSSIAN model was not a universal fit.

**Figure 1:**
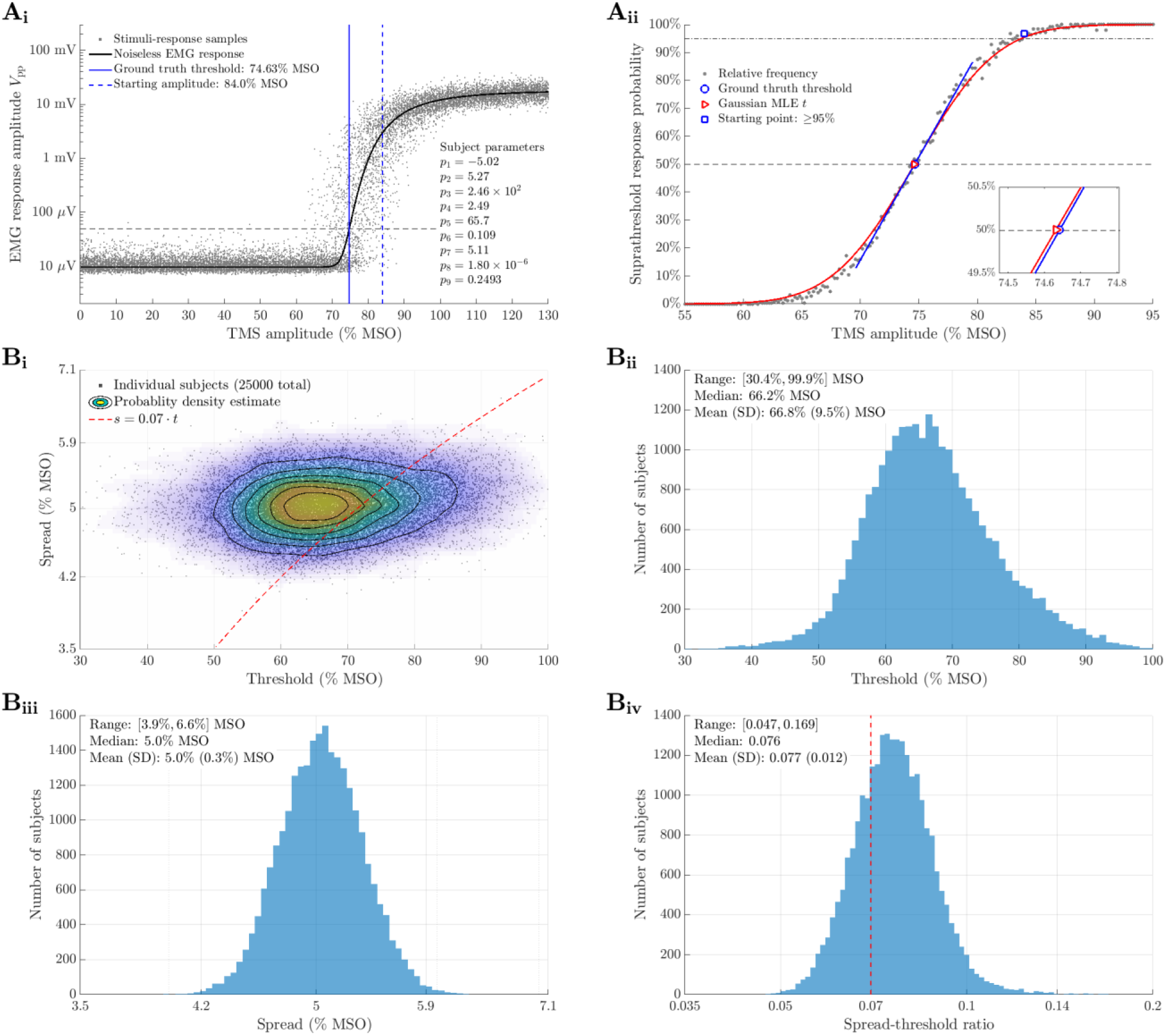
TMS response model. **A**. An example of the TMS sigmoidal recruiting behavior that is mimicked by the model. **i.** In the EMG space, this virtual stimulator provides analog values, which some of the algorithms can utilize. The response model provides uncorrelated EMG response, which are independent from the tested black-box models in the parameter estimators to avoid model-determined advantages. Subsequent responses are not correlated. **ii.** In the probability space, the threshold is determined from the relative frequency of responses by linear regression to find the 50% probability amplitude. The starting amplitude was determined as the minimum amplitude at and above which the response probably was no less than 95%. A Gaussian response model can also be obtained by applying MLE to the stimulation-response pairs in Ai, which fitted well to the relative frequencies for the majority of subject. **B**. The ground truth thresholds and the slope/spread at the threshold are obtained by linear regression for the 25,000 virtual subjects. **i.** Joint distribution of threshold and spread. **ii.-iv.** Distributions of thresholds, spread, and spread-to-threshold ratio. The distributions are normal or log-normal. The nominal 0.07-ratio is shown by red dashed lines.

#### 2.4.2. Simulation routine

The testbed comprised all threshold detection algorithms defined previously and the response model. In order to meet the real conditions of a threshold search, each algorithm was given information from the hotspot detection procedure, i.e., a starting amplitude which was known to be above the threshold [52], [76], [77]. To avoid ambiguity and circular definition between threshold and hotspot [78], the latter was chosen as the amplitude that generated a response for at least 95% of the stimuli (Figure S2). The starting amplitude was on average 8.7% MSO higher than the ground truth threshold (1.13 times threshold). For the algorithms that use specific integer amplitudes (i.e., IFCN relative frequency and MILLS–NITHI methods), the starting amplitude was rounded to 1% MSO.

The responses for subsequent stimuli were provided to the algorithms by the full MEP model. Whereas the analog versions of the stochastic root-finding methods directly processed this continuous information, all other methods quantized the response into the binary representation of above and below threshold first. To examine both their short- and long-term behaviors, the parametric estimation algorithms and stochastic root-finding methods were tested with a maximum of 200 steps each. The absolute and relative errors of each threshold estimate *x*_th,est_ versus the ground truth *x*_th,GT_ were analyzed with

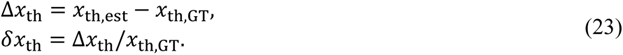

For comparison with previous studies in the literature, the absolute values of the errors |Δ*x*_th_| and |*δx*_th_| were also analyzed, of which |*δx*_th_| was previously referred to as the “relative error” [48], [54]. The results mostly present the relative errors *δx*_th_, as they provide normalized distance of the threshold estimate from the true value expressed as a percentage of the latter. The absolute error Δ*x*_th_ overall had similar distributions and are shown in the supplement.

## 3. Results

### 3.1. IFCN relative-frequency method and MILLS–NITHI two-threshold method

As both methods have been previously established in the field, only the high-level performance is described here, with more details and full statistics provided in the supplement (Tables S1‒S7).

For the IFCN relative-frequency methods (Figures 2 and S3‒S5), the number of stimuli until the method terminated and reported a threshold value for a given subject had a wide distribution. On the lower end, the methods stopped after reducing the stimulation amplitude by two or three steps below the starting amplitude, which could be as few as 15 to 20 stimuli for the five-out-of-ten methods or between 40 and 60 for the ten-out-of-twenty methods. On the other hand, some subjects required a larger number of stimuli, reaching over 100 or even 200 depending on the method. Estimates obtained with fewer stimuli overestimated the threshold due to false early fulfillment of the termination rule, whereas those with large numbers of stimuli leaned towards underestimation due to false continuation at an amplitude with response probability less than 50%. The errors were linearly dependent on the number of stimuli with coefficients of determination (*r*^2^) more than 0.7, and over overall had a positive bias (Figure S5, panels Bi and Bii; e.g., median, mean, and SD of 0.91%, 1.00%, and 1.75%, respectively, for *δx*_th_ of the ten-out-of-twenty method with 1% MSO stepping size). Such linear relationship and bias were inherently determined by the binomial probability distribution to either terminate or continue (Figure S4; see method for calculating the probability in Supplement), consistent with a previous argument against relative-frequency methods [43]. The number of stimuli drastically decreased by approximately 40% and 50% when the stepping size was increased from 1% to 2% MSO and the coarser five-out-of-ten rule was used instead of the ten-out-of-twenty rule for the relative frequency approximation, respectively, although the shape of the distribution was not affected qualitatively. These changes overall decreased the threshold accuracy (i.e., increased median, mean, and standard deviation of errors) as intuitively expected. However, the errors had similar ranges for both ten-out-of-twenty or five-out-of-ten methods with either step sizes. Therefore, it appears ineffective to use smaller step size and/or a larger number of stimuli at a given amplitude to improve the accuracy, as the threshold error of individual subjects does not necessarily decrease but the required number of pulses increases significantly.

**Figure 2:**
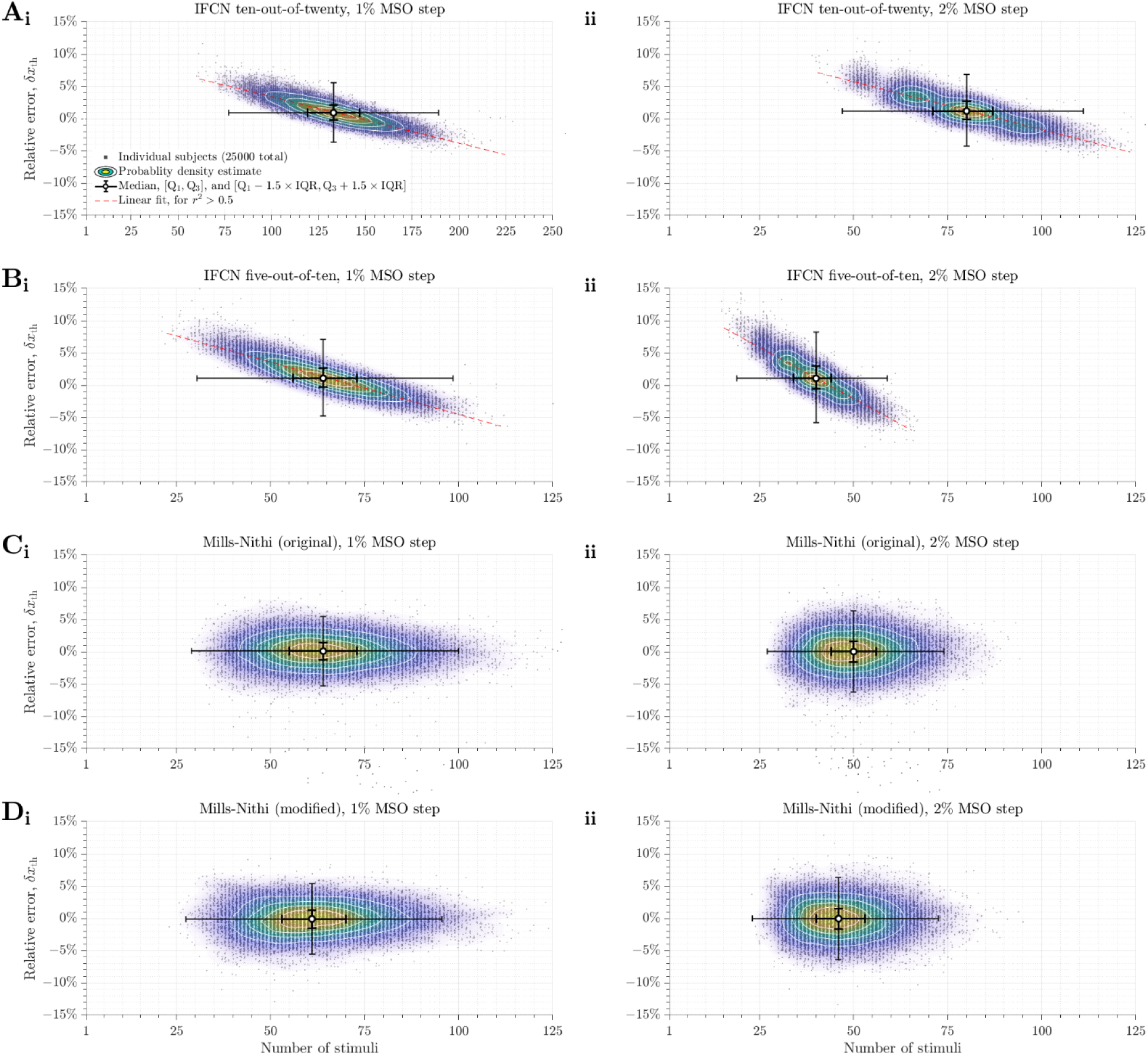
Relative threshold errors of the IFNC relative frequency and MILLS–NITHI methods. The relative errors *δx*_th_ are plotted against the number of stimuli required for all the virtual subjects as dots. Several statistics are shown, with median as white circle, interquartile range between first and third quartiles (Q1 and Q3) as thick error bars, and ranges extending to the lower and upper adjacent values—which are 1.5 times the interquartile range below Q1 and above Q3 and cover approximately ± 2.7 standard deviations for normally-distributed data—as thin error bars. A probability density estimate based on a normal kernel function is also shown with the contour lines. Linear regressions are shown for the IFCN relative-frequency methods as red lines. **A.** IFNC ten-out-of-twenty method. **i.** 1% MSO step size. **ii.** 2% MSO step size. **B.** IFNC five-out-of-ten method. **i.** 1% MSO step size. **ii.** 2% MSO step size. **C.** Original MILLS–NITHI methods. **i.** 1% MSO step size. **ii.** 2% MSO step size. **D.** Modified MILLS–NITHI methods. **i.** 1% MSO step size. **ii.** 2% MSO step size. Note different horizontal scale in panel Ai versus the others.

**Figure 3:**
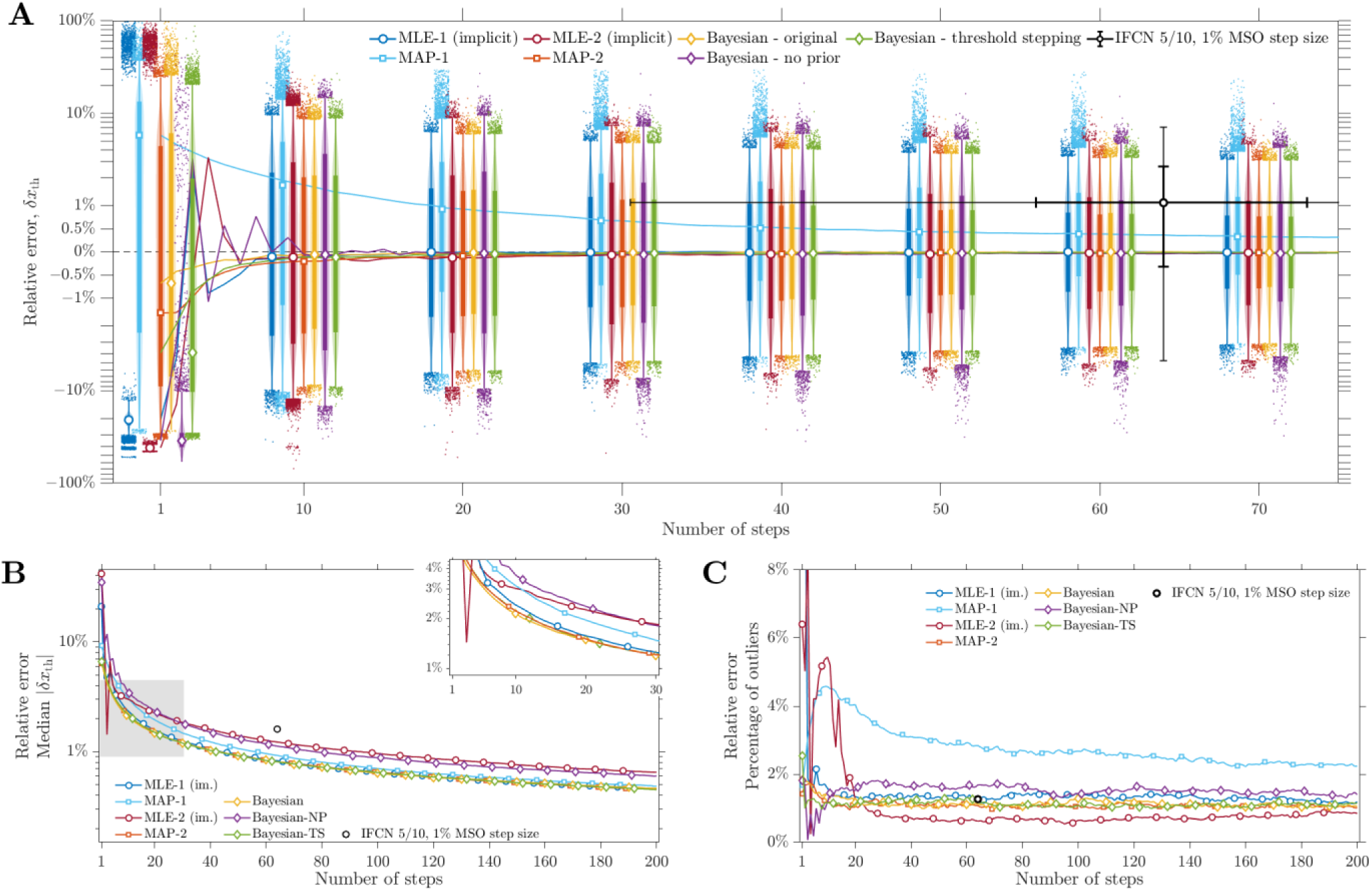
Relative threshold errors of the parametric estimation methods. For MLE methods, results from implicit (im.) maximization of the likelihood function are shown. **A.** Statistics are shown up to the seventieth steps as boxes-and-whiskers at the first step and every ten steps starting from the tenth step, with median as marker, interquartile range as thick vertical line, upper and lower adjacent values as whiskers (thin vertical lines), and outliers as dots. The lines show the median at every step. The boxes-and-whiskers for different methods are slightly offset for clarity. The log-lin-log mixed scale is used to better visualize the errors around zero, with linear scaling in the range of [−1%, 1%] and logarithmic scaling of amplitudes outside this range. The median, interquartile range, and range of adjacent values of the IFCN five-out-of-ten method with 2%MSO step size are shown as a circle and black error bars as in Figure 2Bii for reference. **B.** The median |*δx*_th_| on logarithmic scale, with the inset showing the details in the first 30 steps (the gray area in the main panel). **C.** The percentage of outliers (dots in panel A) outside the upper and lower adjacent values (whiskers in panel A).

The MILLS–NITHI methods (Figures 2, S3 (panels Ci, Cii, Di, and Dii), and S5) had narrow distributions of the number of stimuli, requiring at least 20 stimuli but rarely exceeding 100. The numbers of stimuli for all four variations were on average similar to or smaller than the IFCN five-out-of-ten method with 1% MSO step size but larger than that with 2% MSO step size. In comparison, the IFCN relative-frequency method could be much faster for any individual subject due to its false early termination and/or if the starting point was already close to the threshold (i.e., steep slope near the threshold). The modified MILLS–NITHI methods exploited the additional information obtained from the hotspot search but only reduced the number of stimuli by three or four on average compared to the original method using the same step size. Nevertheless, none of the MILLS–NITHI methods resulted in extremely large numbers of stimuli. Due to the averaging of the upper and lower thresholds, the errors of the MILLS–NITHI methods were independent of the number of stimuli (*r*^2^< 0.01) and, compared to the IFCN relative-frequency methods, had tighter distributions and were centered closer to zero. Compared to the modified methods, the original MILLS–NITHI methods had wider error distributions and many outliers with large errors. Increasing the step size to 2% MSO for either the original or modified method reduced the number of stimuli on average by 14 or 15 steps and narrowed its distribution. Increasing the step size also widened the error distribution slightly.

### 3.2. Parametric estimation

Unlike the relative-frequency methods, the parametric estimation methods generated a threshold estimate and thus an error at every step (relative error δ*x*_th_ in Figure 3A and absolute error Δ*x*_th_ in Figure S6A). The median |δ*x*_th_| (Figure 3B) and the interquartile range (IQR) of δ*x*_th_ characterized how close the errors approach zero and how wide the errors were distributed. The IQR of δ*x*_th_, and therefore the range between the lower and upper adjacent values (which were 1.5×IQR below and above the first and third quartile) as well, correlated well with the median |δ*x*_th_|: the ratio between these two statistics after a few steps was approximately 2 and was very stable (Figure S7). Therefore, the median |δ*x*_th_| captured the distribution of most errors, with the remaining outliers summarized as a percentage of the total number of subjects (Figure 3C). Besides the step-by-step distributions of errors, a few metrics were used to benchmark and compare the performance (Table 4). Estimation speed was quantified by the number of steps for a method to reach the same median |δ*x*_th_| as the IFCN five-out-of-ten method with 1% MSO step size (median of 64 steps and median |δ*x*_th_| of 1.6%). Whereas recent IFCN guidelines [28] recognized that the relative-frequency methods require ten-out-of-twenty trials to generate reliable results, the five-out-of-ten method is still widely used [79]–[82] and therefore chosen as the reference. For accuracy, the median |δ*x*_th_| was compared at 20 and 30 steps, which are respectively considered to produce a sufficiently accurate estimate for all subjects using existing ML-PEST (MLE-1) [41], [48] and commonly used as the upper limit to terminate adaptive thresholding procedures [53], [71], [72].

**Table 4.** Error convergence of parametric estimation and root-finding methods

| Method | Step reaching<br>median $\vert \delta x_{th}\vert $ of 1.6% | Median $\vert \delta x_{th}\vert $<br>at step 20 | Median $\vert \delta x_{th}\vert $<br>at step 30 | Optimal $a_0$<br>(ratio to default value) |
| --- | --- | --- | --- | --- |
| MLE-1 <sup>a</sup> | 20 (21) | 1.57% (1.62%) | 1.25% (1.26%) | — |
| MAP-1 | 27 | 1.96% | 1.48% |  |
| MLE-2 <sup>a</sup> | 40 (39) | 2.26% (2.27%) | 1.85% (1.83%) |  |
| MAP-2 | 18 | 1.506% | 1.21% |  |
| BAYESIAN | 18 | 1.48% | 1.198% | — |
| BAYESIAN-NP | 37 | 2.37% | 1.81% |  |
| BAYESIAN-TS | 18 | 1.50% | 1.196% |  |
| ACS1-H <sup>b</sup> | 24 (18) | 1.69% (1.509%) | 1.46% (1.31%) | ×1.6 |
| ACS2-H | 28 | 1.77% | 1.56% | — |
| DCS1-H <sup>c</sup> | 17 (16) | 1.450% (1.446%) | 1.17% (1.18%) | ×0.79 |
| ACS1-HA <sup>c</sup> | 18 (18) | 1.517% (1.512%) | 1.280% (1.30%) | ×0.79 |
| ACS2-HA | 18 | 1.521% | 1.27% | — |
| DCS1-HA <sup>b</sup> | 22 (19) | 1.67% (1.54%) | 1.37% (1.24%) | ×0.63 |
| ACS1-GA <sup>c</sup> | 20 (20) | 1.59% (1.584%) | 1.45% (1.47%) | ×0.79 |
| ACS2-GA | 19 | 1.581% | 1.439% | — |
| DCS1-GA <sup>c</sup> | 22 (21) | 1.68% (1.62%) | 1.43% (1.444%) | ×0.63 |
| ACS1-GHA <sup>b</sup> | 19 (18) | 1.53% (1.514%) | 1.283% (1.29%) | ×0.63 |

<sup>a</sup> Results obtained with direct maximization of the likelihood followed by those using explicit evaluation in parentheses.
<sup>b</sup> Results obtained with default $a_0$ followed by those with optimal $a_0$ for lowest median $|\delta x_{th}|$ at both step 20 and 30 in parentheses.
<sup>c</sup> Results obtained with default $a_0$ followed by those with optimal $a_0$ for lowest median $|\delta x_{th}|$ at step 20 in parentheses, with default $a_0$ being optimal for lowest median $|\delta x_{th}|$ at step 30.

The basic MLE method [48], which used a single-parameter MLE for threshold estimation and stepping (MLE-1, Figure 3, dark blue lines with circle markers), reached the same error level as the reference IFCN five-out-of-ten method with only 20 steps (Table 4), and the error further decreased by about 20% at 30 steps. Evaluating the likelihood function explicitly had little effect on the threshold estimation performance (Table 4, results in parentheses, and Figure S8) and computational runtime (Figure S9) compared to implicit maximization. Introducing prior information of the parameters of the population (MAP-1, Figure 3, light blue lines with square markers) did not improve the performance for the single-parameter estimation. On the contrary, with the empirical relationship *s* = 0.07 ⋅ *t* from another population imposed on the parameter distribution of the virtual population in this study, MAP-1 could not fully utilize the prior distribution. Its performance was considerably worse than MLE-1 as the parameters of some subjects in our model were far from such a 0.07 relationship (Figure 1Biv).

Compared to MLE-1, the two-parameter MLE method (MLE-2, Figure 3, dark red lines with circle markers) had a slower convergence and reached the median error level of the IFCN reference with more (40) pulses. The reduced speed of convergence reflected the exploration of the full parameter space: the likelihood function initially had a large plateau region, which reduced in size as more stimulation–response pairs were recorded. Only after a sufficient number of steps did the likelihood plateau narrow to generate a clear optimal estimate. Similar to MLE-1, the method for maximizing the likelihood function had little effect on performance for MLE-2 (Table 4, results in parentheses, and Figure S8). However, the explicit evaluation with two parameters resulted in a much higher computational runtime per step that was two order-of-magnitudes slower (Figure S9). Using a two-parameter MAP estimator for the threshold (MAP-2, Figure 3, orange lines with square markers) significantly improved the performance of MLE-2, reaching the error level of the IFCN five-out-of-ten method with only 18 steps, and had lower median |δ*x*_th_| compared to either MLE-1 or MAP-1. As the spread parameter was no longer fixed to the threshold, MAP-2 could match the wide range of behaviors of the subjects in our model and improved the threshold estimate for all subjects. The prior information significantly influenced the likelihood landscape, rendering one parameter pair (*t*, *s*) in the otherwise flat maximum plateau unique. The computational runtime of MAP-1 and MAP-2, which used implicit evaluation, were slightly faster their implicit MLE counterparts (Figure S9).

The original KONTSEVICH–TYLER method (BAYESIAN, Figure 3, yellow lines with diamond markers) had the least bias in error distribution (Figure 3A), reached the IFCN five-out-of-ten method’s median |δ*x*_th_| quickly at step 18 and had faster convergence for small number (< 20) of stimuli (Figure 3B). The BAYESIAN method achieved more accurate estimation early on by considering the entire function space of the probability density functions. Whereas ML and MAP estimators evaluated the underlying likelihood function in every step anew based on the same key equation, the BAYESIAN approach added information into the explicitly evaluated and stored distributions from the last iteration. However, such advantages disappeared with more pulses as the point estimator methods gathered sufficient information. Removing the prior population information (BAYESIAN-NP, Figure 3, purple lines with diamond markers) resulted in slower convergence and performance comparable to that of MLE-2 with a similar slow down comparing MAP-2 to MLE-2, showing that the prior population information narrowed the distribution of parameters in the probability space alike for the point and distribution estimators. Utilizing the threshold estimate for stepping (BAYESIAN-TS, Figure 3, green lines with diamond markers) had similar performance compared to the original method of minimizing the expected entropy, demonstrating that the threshold was at or very close to the stimulation amplitude that provided the maximum information gain. This provided further validation for choosing the threshold estimate as the next stimulation amplitude for the MLE and MAP methods, despite the difference of how the estimate was obtained—for the BAYESIAN methods, the threshold was estimated as the expected value based on its probability distribution (21), whereas for MLE and MAP methods the estimation came from maximizing the likelihood functions (13)‒(14). The BAYESIAN methods were the most computationally costly and slowest (Figure S9), with runtime around four seconds per step and more than two order-of-magnitudes slower than the implicitly evaluated MLE/MAP methods. This was mostly due to updating of the entire probability distribution (16), as the runtime without the entropy calculation in BAYESIAN-TS was only slightly reduced and still on the same order-of-magnitude. The BAYESIAN methods were also highly sensitive to rounding-error propagation and sampling of the parameter space, which was a disadvantage compared to MLE and MAP methods.

### 3.3. Stochastic root-finding methods

The parameter-free stochastic root-finding methods were based on the threshold definition only and therefore had the important advantage over the parametric estimation approaches, which depended on a potentially biased or inaccurate model. Their simple stepping rule was also much faster to compute, with runtime per step on the millisecond scale. Like the parametric methods, the root-finding methods also generated a threshold estimate and an error at every step (relative error δ*x*_th_ in Figures 4, S13A, and S15A, and absolute error Δ*x*_th_ in Figures S10, S13B, and S15B). The ratio between the IQR of δ*x*_th_ and the median |δ*x*_th_| (Figures S11 and S14) was also centered on 2 after a few steps, but showed larger variations compared to those of parametric estimation methods.

**Figure 4:**
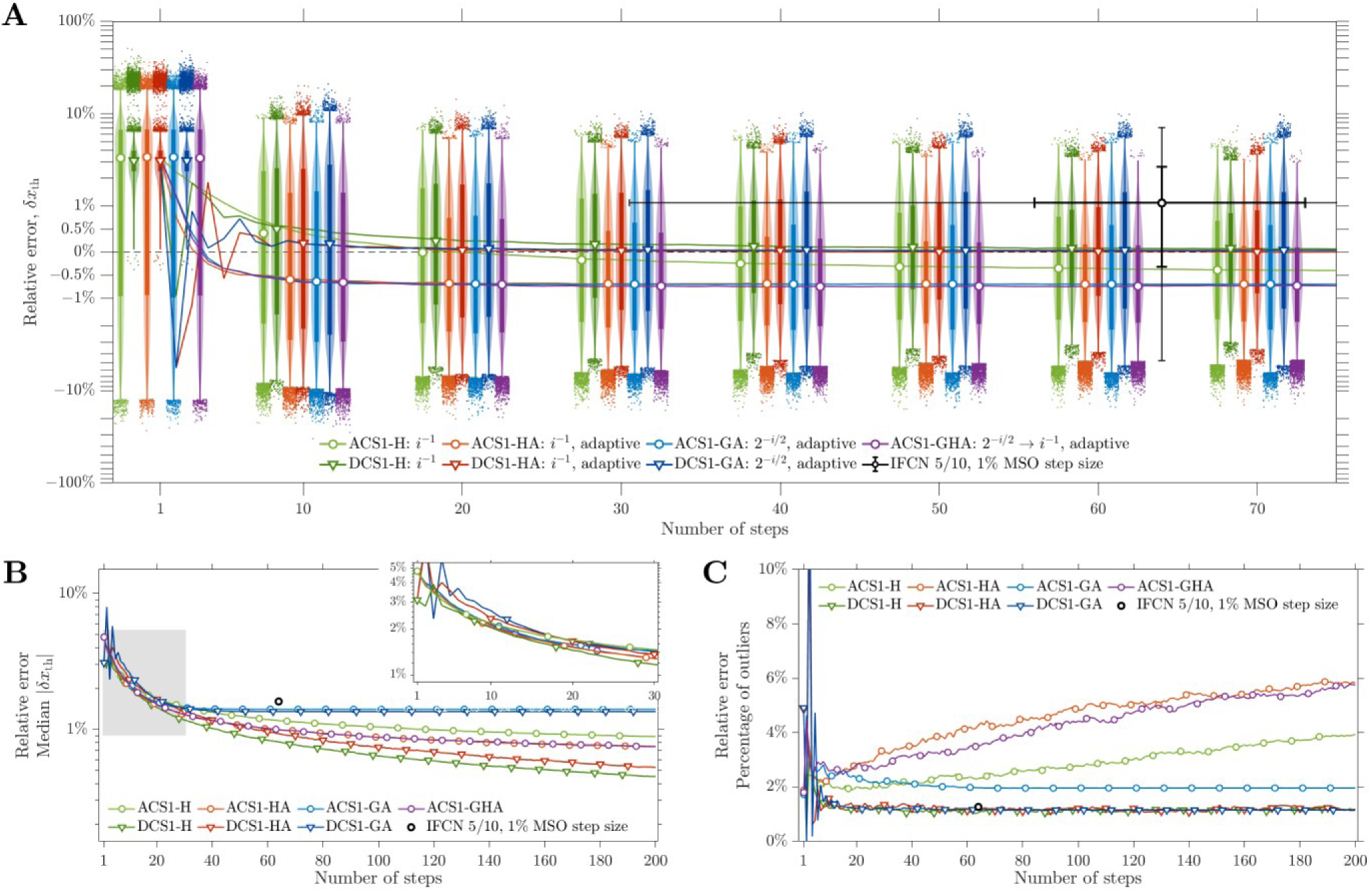
Relative threshold errors of root-finding methods using first-order control sequences with default initial step size *a*_0_. Similar to Figure 3.

The error distributions of all first-order analog methods (Figure 4, lines with circle markers) became negatively biased after a certain number of steps, i.e., the median was less than zero, the absolute values of the first quartile and lower adjacent value were larger than the third quartile and upper adjacent value, respectively, and there were more negative than positive outliers (Figures 4A and S10A). ACS1-H took about 15 and 50 steps for the bias to become negative and plateau, respectively, whereas the process for the other methods was much quicker (within ten steps). The bias originated from the asymmetry of the log-transformed MEP distribution near the threshold [61, Fig. 2]. The 50 μV threshold (which is the median of the distribution according to the probability definition of stimulation threshold) was smaller than the average of the right-skewed distribution and near the EMG noise floor (around 10 μV). Therefore, subthreshold responses generated a smaller Δ*y* to increase a subthreshold stimulation amplitude towards the threshold. In contrast, suprathreshold MEPs can be orders-of-magnitudes larger than the threshold, resulting in large Δ*y* that decreased the stimulation amplitude much lower than the threshold, especially during the early steps. Three of the analog root-finding methods (ACS1-HA, ACS1-GA, and ACS1-GHA) with default parameters reached the accuracy level of the IFCN five-out-of-ten method with 1% MSO step size with only 20 steps or less (Figures 4B and Table 4), whereas ACS1-H needed 24 steps.

The non-adaptive ACS1-H (Figure 4, green lines) had a monotonically declining median |δ*x*_th_| for the entire range of steps. The percentage of outliers, however, increased with the steps, showing that the error did not improve for some subjects, which became a larger proportion of the total population as the errors of the other subjects in the population became more tightly distributed. The adaptive ACS1-HA (Figure 4, orange lines) outperformed its non-adaptive counterpart by reaching a lower median |δ*x*_th_| faster, but with distributions of δ*x*_th_ being more negatively biased and having larger proportion of outliers. The ACS1-GA method (Figure 4, blue lines) with higher rate of convergence had a rapid exponential decay of the control sequence and did not further head for the threshold as soon as *a*_*i*_ was too small for a notable influence. It indeed plateaued earlier, at around 40 steps and at a higher median |δ*x*_th_| of 1.4%, albeit with a smaller portion of outliers as its distribution did not continue to tighten. In comparison, ACS1-GHA (Figure 4, purple lines) switched from fast geometric convergence that provided the rough estimate to a. s. harmonic convergence and performed very similarly compared to ACS1-HA in terms of median |δ*x*_th_| and percentage of outliers.

Besides the convergence of the control sequence *a*_*i*_, the root-finding methods’ performance was also determined by the initial value *a*_0_of the sequence. The default value of *a*_0_(11) was chosen to approximate the typical slope of the recruitment curve but its definition was inevitably empirical and potentially suboptimal. Examining the performance as a function of the initial step size indeed revealed that this was the case for some of the root-finding methods. The change from overshoot and oscillatory behavior for large *a*_0_ towards lengthy one-sided convergence for smaller numbers was clearly visible (Figure S12, panels Ai–Gi). Some methods did not have one single optimal *a*_0_, however, as the minimum of the median |δ*x*_th_| at different steps (Figure S12, panels Aii–Gii, black dashed line and dots) may not be obtained with the same *a*_0_. Instead of evaluating asymptotic behavior, we chose the optimal *a*_0_ that minimized the error at step 20 for which the default was not optimal and the performance indeed improved (Table 4, results in parentheses, and Figure S13). The optimal *a*_0_ for most methods was close to the default value, showing that using the slope of typical input–output curves (11) indeed provided an appropriate *a*_0_. For ACS1-H, the optimal *a*_0_also resulted in a negative bias of error distribution in just a few steps, similar to the other analog versions.

When the default *a*_0_ was used, the digital versions of the root-finding methods (Figure 4, darker lines with triangle markers) could asymptotically perform either better than (DCS1-H and DCS1-HA) or very similar to (DCS1-GA) their analog counterpart in terms of median |δ*x*_th_| (Figure 4B). As the root-finding approached the target, the difference between the actual and target responses became very small (|*a*_*i*_Δ*y*_*i*_| ≪ 1). The step towards the target |Δ*x*_*i*_| was therefore smaller for the analog version (|*a*_*i*_Δ*y*_*i*_|) compared to the digital version (|*a*_*i*_*δy*_*i*_| = *a*_*i*_) and thus approached the target slower than the latter. It should be further noticed that all digital versions had mostly positive bias for δ*x*_th_ that decayed to zero, with the decay being especially quick for the two adaptive methods. The lack of asymptotic bias in contrast to the analog versions was due to the sign function of the digital versions equalizing the stepping towards threshold in both directions. With optimal *a*_0_, the digital versions had a slower decay of the positive bias of the error distribution compared to those for default *a*_0_ (Figure S13Ai) but had improved convergence of |δ*x*_th_| (Figure S13Aii).

The methods with second-order control sequences (Figure S15, lighter lines with square markers) could perform either slightly better (ACS2-HA and ACS2-GA) or worse (ACS2-H) than their first-order counterpart when the default parameters were used. Their performance depended on both the initial step size *a*_0_and the second-order weight *w* and the two parameters interacted as demonstrated with ACS2-HA (Figures S16 and S17). Large/positive *w* sped up the approach towards the threshold, but generated oscillations similar to large *a*_0_. Small/negative *w*, like small *a*_0_, however, reduced the speed of convergence. The second-order weight could compensate a suboptimal *a*_0_ to generate an optimal approach, and vice versa. Compared to the slope-like *a*_0_(10), an optimal *w* was difficult to determine without empirical information or performing extensive parameter sweeps. Therefore, the addition of a second-order term to the root-finding would be less preferable than optimizing the initial step size for the first-order methods.

## 4. Discussion

We examined a variety of existing and new TMS thresholding methods and compared their performance in a virtual population of 25,000 subjects. The methods included eight based on relative frequencies using either a single threshold (IFCN relative-frequency methods) or two thresholds (MILLS–NITHI methods) and either 1% MSO or 2% MSO step size, seven parametric estimation methods, of which four used different combinations of single- or two-parameter ML or MAP estimation and three were based on BAYESIAN estimation that updated the probability density function itself, and ten stochastic root-finding methods utilizing control sequences with different convergence and stepping rules.

Although there were some trade-offs between different methods, several threshold estimation methods can be considered to have overall better performance within their class or even across all methods (Figures 5 and S18). All relative-frequency methods had similar error performance and should be avoided due to the large number of pulses required and the presence of inherent error bias or large error outliers (see Figure S5). The two-parameter MAP estimation demonstrated the best performance among the point estimators considering the speed of convergence, median error, and computational complexity. The KONTSEVICH–TYLER BAYESIAN estimation performed even better regarding the first two aspects, but with increased computational cost, which might mostly matter if implemented on embedded microcontrollers. The best stochastic root-finding methods using analog control sequences was ACS1-HA, whereas DCS1-H had the strongest performance among both analog and digital versions. Unsurprisingly, the best-performing novel methods as well as the previous single-parameter MLE all outperformed the reference IFCN five-out-of-ten method (median 64 steps and 1.6% median |δ*x*_th_|), requiring much fewer steps (within 20 steps, with the novel methods a few steps faster than the single-parameter MLE) to reach the same accuracy (Figure 5B, top). Compared to MLE-1, the best-performing novel methods also demonstrated better accuracy at 20 steps (Figures 5A and S18), which is a typical number used to terminate the single-parameter MLE methods [41], [48], for example in ATH-tool [72].

**Figure 5:**
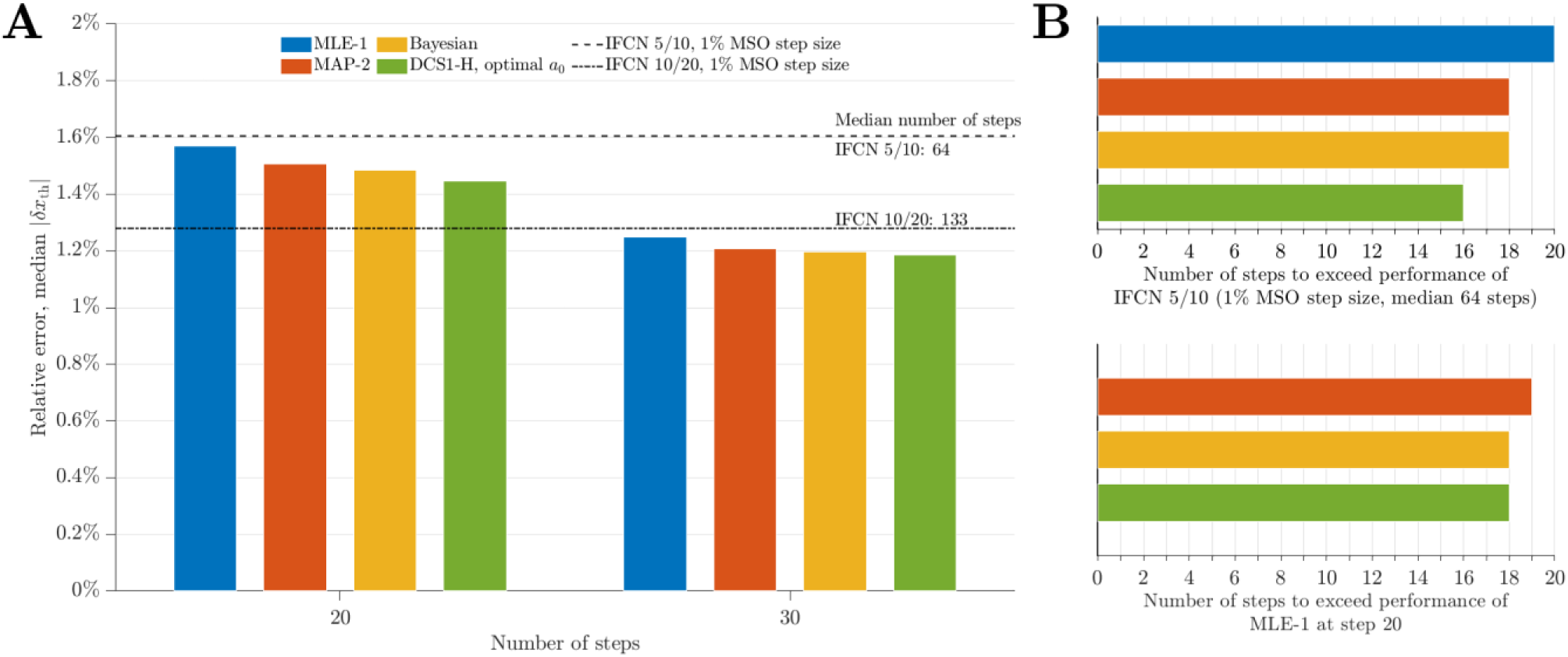
Summary of best performing methods. Selected statistics of the best performing methods, with full statistics shown in Figure S18. The best parametric methods are two-parameter MAP estimation (MAP-2) and BAYESIAN estimation, with the single-parameter MLE method (MLE-1) given for comparison. The best stochastic root-finding methods is DCS1-H, with optimal *a*_0_shown. The MAP-2 and BAYESIAN methods utilize prior information about the threshold distribution. **A.** Median |*δx*_th_| at steps 20 and 30. The dashed and dotted lines shows for reference the IFCN five-out-of-ten and ten-out-of-twenty methods with 1% MSO step size, respectively. **B.** The number of steps needed for a given method to outperform the reference IFCN five-out-of-ten method (top) and the single-parameter MLE at 20 steps (bottom).

Despite their wide range of behavior and difference in performance, all methods performed remarkably accurate in terms of threshold error on the population level. Either *δx*_th_ and Δ*x*_th_ (or |*δx*_th_| and |Δ*x*_th_|) had mean or median on the level of 1% and 1% MSO, respectively. In comparison, the pulse amplitude variability of TMS machines themselves have exactness (deviation from nominal value) and precision (variation between pulses) in comparable ranges (1% to 3%) depending on the device [83]. The errors of individual subjects, however, had a large spread, and their relative error may not fall within the ±5% criteria for safety [12], [42] for either relative-frequency methods (Figure S5) or more advanced estimation methods (Figures 3A, 4A, and S13Ai). For some methods, the estimation may not improve with increasing number of TMS pulses for certain subjects (e.g., ACS1-H) or the entire population (e.g., ACS1-GA) after a certain number of steps.

We analyzed both the errors and their absolute values, and provided detailed visualization and descriptive statistics of the entire distribution. In contrast, many previous studies quantified the absolute values of the relative error (|*δx*_th_|, simply referring to it as the relative error) and/or presented mean and standard deviation only [48], [54]. The use of absolute values for errors provided the convenience to compare a positive metric that decreased with stepping and allowed the use of logarithmic scale for visualization. Whereas the median of the absolute values can capture the spread of the errors very well—our results showed that the ratio between the IQR of the errors and the median of the errors’ absolute value was quite consistently around two for the methods examined—the absolute value of errors, however, could not capture the bias of the estimated thresholds or its symmetry around zero. Similarly, the mean and standard deviation did not capture outliers and skewness of the distributions, which were all important characteristics of the thresholding methods. For example, the relative-frequency methods had inherent bias regardless of stepping size or the number of stimuli at a given amplitude, and the original MILLS–NITHI methods had a large number of outliers with negative error, contradicting assertions about its reliability [42].

Our implementation of the various methods all started from a suprathreshold simulation amplitude with a minimum 95% response probability to avoid ambiguity and circular definition of hot-spot and threshold. This, however, is often not the case in practice. A lower starting amplitude could be used to accelerate the threshold search for some methods, especially for subjects with shallow input–output curves, whereas a default stimulator output is sometimes used as starting point for all subjects. The stimulation amplitudes of stochastic root-finding methods are constrained by previous pulses, and different starting points change the error convergence and require a different optimal initial step size *a*_0_(results not shown). In contrast, the starting point does not affect the performance of parametric estimation methods on the population level, e.g., using a fixed 50% MSO amplitude (results not shown), as their stimulation amplitudes determined by the likelihood maximization are not constrained and can freely “jump” arbitrary distances from pulse to pulse. However, both of these situations risk convergence errors when a positive response is observed for a stimulator output well below the true threshold, especially early during the procedure [52], [53].

Existing estimation methods previously were tested using virtual subjects that had the same response model underlying the parameter estimation methods [48]. Such tests of estimators against their own estimation model can be misleading due to statistical bias. We therefore used an independently derived MEP model from the literature, which was previously trained to the statistics of experimental data. The response model used here, although more sophisticated than most other approaches, may still have limitations in representing all features and statistical properties of stimulation-evoked EMG responses due to the limited data on which it was based [32], [84]–[86]. In addition, complex temporal effects, such as inter-pulse correlation caused by stimulation-induced short-term plasticity and also stimulation-independent excitability fluctuations [45] may become important, particularly for short inter-pulse intervals of only a few seconds used in some labs. Accurate knowledge of the physiologic system, however, is essential for the thresholding problem. This does not only influence the design of a realistic testbed, but also directly determines, for example, how well the GAUSSIAN model used in all parameter estimators suits the requirements or if another model, such as logistic distribution [40], [50], is needed in order to avoid bias.

The performance of some estimation methods demonstrated substantial improvement over existing methods. For example, the two-parameter MAP and KONTSEVICH–TYLER BAYESIAN estimation methods all outperformed the single-parameter MLE methods. The improvement in accuracy for the BAYESIAN method, however, comes at a cost of increased runtime per step and larger memory requirement. While these costs may still be challenging for many microcontrollers depending on the specific application and potential implementation as an embedded version on a TMS device, they are readily manageable on personal computers which many recent TMS machines anyway include. The BAYESIAN method could become more appealing with improved implementation efficiency, for example by reducing the range and/or resolution of parameter space. All parametric estimators, however, had the potential weakness that the underlying model could be incorrect. Some of the parametric methods further relied on accurate prior information of the model parameters (threshold and spread in the GAUSSIAN model) to achieve the presented accuracy and improvement over other methods. Such prior information on the population level could require significant effort to collect or may be completely unavailable.

The newly introduced family of non-parametric stochastic root-finding methods overall had similar or even lower errors compared to the parametric estimation methods. Comparing the best performing methods between the two classes, DCS1-H was superior in terms of both speed and accuracy compared to the two-parameter MAP estimation and KONTSEVICH–TYLER BAYESIAN methods, which both required prior information. Furthermore, stochastic root-finding methods did not rely on potentially incorrect models and thus could work for subjects whose MEP response curves did not follow the GAUSSIAN model. The rule for choosing and updating the step size *a*_*i*_ was also simple enough to be performed by hand or a calculator and would not necessarily rely on *installed* software (e.g., [48], [72]). Hence, the root-finding methods are a promising addition to the available threshold methods that motivate further studies and improvements. To enable the TMS community to use this method, we implemented DCS1-H as an easy-to-use web-based tool SAMT (Stochastic Approximator of Motor Threshold [87]) using HTML/JavaScript, which is accessible via web browsers on desktop computers and mobile devices.

## 5. Conclusion

This work analyzed three classes and a total of 25 variations of TMS thresholding methods and compared their performance, including their accuracy and speed. As pointed out in previous critiques, the relative-frequency methods, such as five-out-of-ten, were far from optimal due to their large number of required stimuli, wide distribution of errors, and inherent error bias. Their accuracy was outperformed by parametric estimators and stochastic root-finding methods with a fraction of the stimuli. As the response model underlying parametric estimation methods did not always fit well with the virtual subject data, some parametric methods performed poorly for a small subset of subjects in the population. Including all model parameters in the estimation and incorporating prior information boosted performance. Stochastic root-finding methods were comparable or better in performance than parametric estimation methods, were much simpler to implement, and did not rely on an underlying model.

## Code availability statement

The MATLAB code of the thresholding methods is available at [67] (DOI: <u>10.5281/zenodo.6483601</u>). SAMT is available to the community online at [87]: https://tms-samt.github.io.

## Acknowledgments

Research reported in this publication was supported by the U.S.A. National Institutes of Health under Award Numbers R01NS117405 and RF1MH124943. The content is solely the responsibility of the authors and does not necessarily represent the official views of the funding agency. The authors thank Dr. Philip Whiting and Dr. Lari M. Koponen for helpful discussions. Computational support was provided by the Duke Compute Cluster.

S. M. G. conceived, supervised, and secured funding for the study, and developed the theoretical framework and original simulation code. B.W. reconstructed, improved, and optimized the computational framework, performed the simulations, data analysis, and visualization. A.V.P. contributed concepts, supervised, and secured funding and computational resources for the study. B. W. and S. M. G. wrote the manuscript, and all authors revised, commented on, and approved the final version of the manuscript. Preliminary results of this study were presented at the 5th International Brain Stimulation Meeting [88] (February, 2023, Lisbon, Portugal).

## Conflicts of interests

S. M. Goetz and A. V. Peterchev are inventors on patents and patent applications on TMS technology. S. M. Goetz has previously received research funding from Magstim as well as royalties from Rogue Research. A. V. Peterchev has equity options in and serves on the scientific advisory board of Ampa and has received research funding, travel support, patent royalties, consulting fees, equipment loans, hardware donations, and/or patent application support from Rogue Research, Magstim, MagVenture, Neuronetics, BTL Industries, Magnetic Tides, Ampa, and Soterix Medical. B. Wang declares no relevant conflict of interest.

## Appendix

### Response model parameters

The individual recruitment parameters *p*_1,*j*_ to *p*_5,*j*_ followed normal and lognormal distributions [61]:

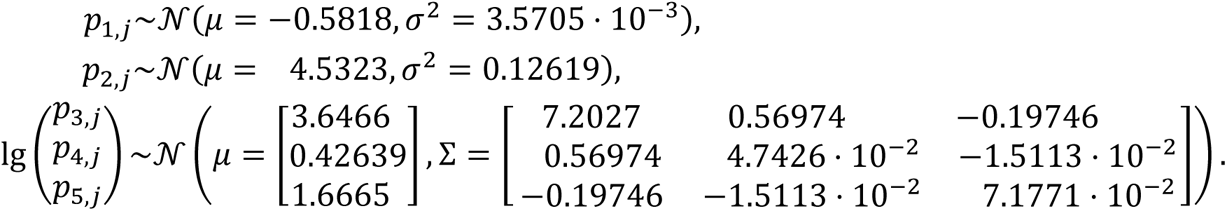

The intra-individual variability sources *ϵ*_xadd,*ij*_ and *ϵ*_ymult,*ij*_ followed normal distributions with subject-specific standard deviation at each step *i*:

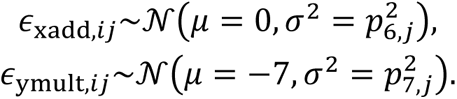

in which the standard deviations for subjects *j* followed lognormal distributions:

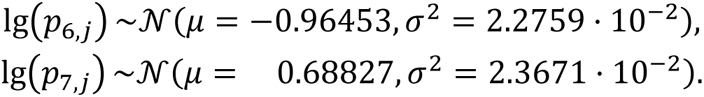

The additive noise *ϵ*_yadd,*ij*_ followed a generalized extreme-value (GEV) distribution, with subject-specific parameters

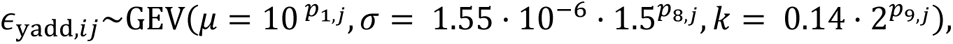

where *p*_8,*j*_ and *p*_9,*j*_ were independent normal distributions 𝒩(0,1), and the GEV distribution had a probability density function of

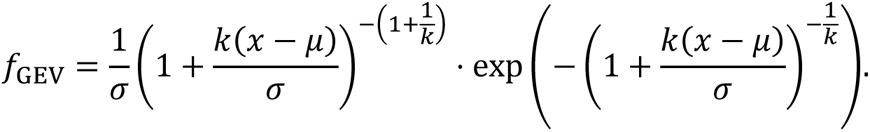

The noiseless model had noise parameters *ϵ*_xadd,*ij*_ and *ϵ*_ymult,*ij*_ set to zero and *ϵ*_yadd,*ij*_ = 10 ^*p*1,*j*^, which approached the mean of the GEV for the parameters *p*_1,*j*_, *p*_8,*j*_, and *p*_9,*j*_.

### Distribution of threshold and spread

The threshold *t* and spread *s* (Figure 1B) was fitted by a joint distribution

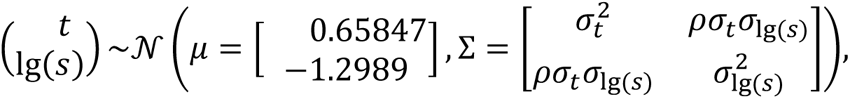

with *σ*_*t*_ = 8.9703 · 10^−2^, *σ*_lg(*s*)_ = 2.7576 · 10^−2^, and the correlation coefficient *ρ* = 0.130. This distribution served as *p*(**θ**) for the MAP methods and *p*_0_(**θ**) for the BAYESIAN methods.

## Supplement

### Numeric calculation of error distribution of IFCN relative-frequency methods

The distribution of the absolute errors of the IFCN relative-frequency methods can be numerically calculated using probability. The number of stimuli for the procedure is *N* (even, e.g., 10), and at the *l*^th^ stimulation amplitude *x*_(*l*)_, the suprathreshold response probability *p*_(*l*)_ follows the Gaussian distribution (12). Here, the stimulation amplitude is relative to the threshold, so the *t* parameter is set to zero in (12). The probability that the procedure stops or continues after *k*_(*l*)_ stimuli are then given based on binary distributions

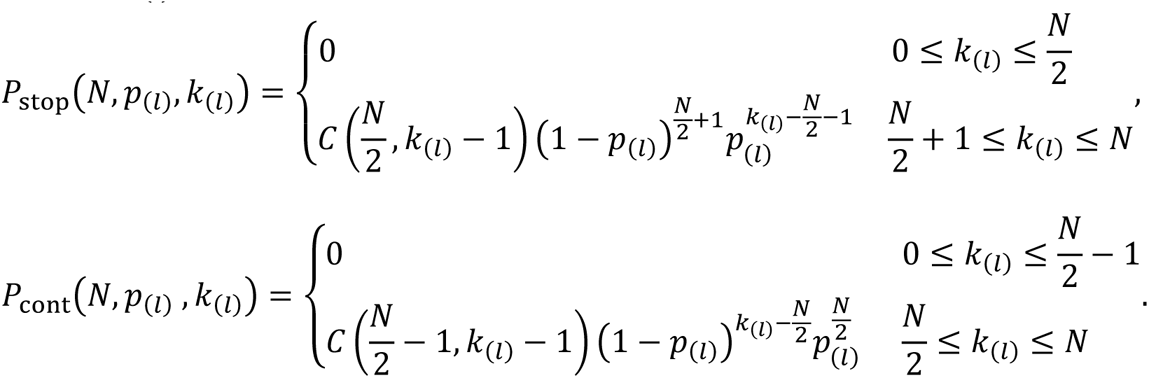

Here, *C*(*n*, *k*) is the number of *k*-combinations for a set of *n* elements. If the procedure continues, the stimulation amplitude is reduced to *x*_(*l*+1)_ = *x*_(*l*)_ − *δx* with *δx* being either 1% or 2% MSO, and the response probability is updated to *p*_(*l*+1)_ by (12) accordingly. This iterative procedure starts with *x*_(1)_ that corresponds to *p*_(1)_ = 0.95 for the given spread *s* and a total probability of one. The probability for the procedure to terminate at a specific amplitude *x*_(*l*)_ for given number of total stimuli *L* can then be calculated as

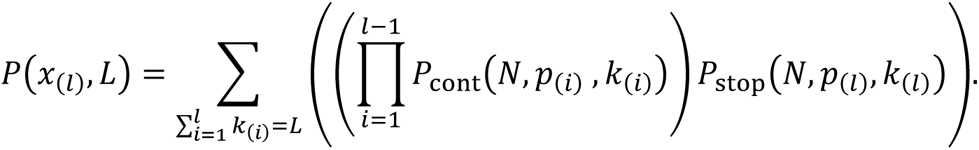

The summation is over all combinations of stimuli numbers *k*_(*i*)_ with non-zero *P*_cont_ or *P*_stop_ that result in a total number *L*.

A recursive depth-first-search was used for the calculation. Besides the natural termination due to failing the relative-frequency criteria, the procedure also was also terminated at points where *p*_(*l*)_ < 0.05, *P*_cont_ < 10^−6^, or the total probability to continue is less than 10^−10^to limit the search depth and speed up calculation. To account for subjects with different response curves (12), i.e., different *s*, *P*(*x*_(*l*)_, *L*) was calculated for a range of spread parameter, and the final distribution was a weighted sum with weights based on the probability distribution of *s* (see Appendix).

## Supplementary tables

**Table S1.** Statistics of number of steps, absolute errors, and relative errors of IFCN and MILLS–NITHI methods. (related to Figures 2, S3, and S5)

| Group number | Method | Step size (% MSO) | Number of stimuli | | Relative error $\delta x_{\text{th}}$ | | | | $\vert \delta x_{\text{th}}\vert $ | Absolute error $\Delta x_{\text{th}}$ (% MSO) | | | | $\vert \Delta x_{\text{th}}\vert $ (% MSO) |
| --- | --- | --- | --- | --- | --- | --- | --- | --- | --- | --- | --- | --- | --- | --- |
| | | | Mean±S.D. | Median | Mean±S.D. | Median | Slope | $r^2$ | Median | Mean±S.D. | Median | Slope | $r^2$ | Median |
| 1 | IFCN, ten-out-of-twenty | 1 | 133.0±20.7 | 133 | 1.00%±1.75% | 0.91% | −0.072% | 0.73 | 1.28% | 0.66±1.13 | 0.60 | −0.047 | 0.74 | 0.84 |
| 2 | IFCN, ten-out-of-twenty | 2 | 79.7±12.1 | 80 | 1.31%±2.11% | 1.17% | −0.15% | 0.72 | 1.58% | 0.85±1.37 | 0.78 | −0.097 | 0.73 | 1.06 |
| 3 | IFCN, five-out-of-ten | 1 | 64.5±12.3 | 64 | 1.23%±2.28% | 1.07% | −0.16% | 0.78 | 1.61% | 0.80±1.47 | 0.72 | −0.11 | 0.79 | 1.07 |
| 4 | IFCN, five-out-of-ten | 2 | 39.6± 7.2 | 40 | 1.22%±2.66% | 1.06% | −0.32% | 0.75 | 1.85% | 0.80±1.73 | 0.71 | −0.21 | 0.76 | 1.24 |
| 5 | MILLS–NITHI, original | 1 | 64.6±13.4 | 64 | 0.03%±2.45% | 0.09% | −0.015% | 0.006 | 1.35% | 0.02±1.61 | 0.06 | −0.01 | 0.007 | 0.89 |
| 6 | MILLS–NITHI, original | 2 | 50.5± 9.3 | 50 | −0.04%±2.69% | 0.02% | −0.003% | 10<sup>−4</sup> | 1.57% | −0.03±1.78 | 0.01 | −0.002 | 10<sup>−4</sup> | 1.05 |
| 7 | MILLS–NITHI, modified | 1 | 62.0±13.2 | 61 | −0.11%±2.10% | −0.10% | 0.004% | 10<sup>−4</sup> | 1.38% | −0.07±1.36 | −0.07 | 0.003 | 10<sup>−4</sup> | 0.91 |
| 8 | MILLS–NITHI, modified | 2 | 47.0± 9.2 | 46 | −0.06%±2.41% | −0.05% | −0.012% | 0.002 | 1.59% | −0.04 ±1.56 | −0.03 | −0.008 | 0.002 | 1.05 |

**Table S2.** Analysis of variance (ANOVA) of number of stimuli of IFCN and MILLS–NITHI methods. (related to Table S1)

| Source | SS | df | MS | F | Prob>F |
| --- | --- | --- | --- | --- | --- |
| Groups | $1.495 \times 10^8$ | 7 | $2.135 \times 10^7$ | $1.308 \times 10^5$ | 0 |
| Error | $3.265 \times 10^7$ | 199992 | 163.2 | | |
| Total | $1.821 \times 10^8$ | 199999 | | | |

**Table S3.** ANOVA of relative errors of IFCN and MILLS–NITHI methods. (related to Table S1)

| Source | SS | df | MS | F | Prob>F |
| --- | --- | --- | --- | --- | --- |
| Groups | $7.950 \times 10^4$ | 7 | $1.136 \times 10^3$ | $2.171 \times 10^3$ | 0 |
| Error | $1.046 \times 10^6$ | 199992 | 5.231 | | |
| Total | $1.126 \times 10^6$ | 199999 | 0 | | |

**Table S4.** ANOVA of absolute errors of IFCN and MILLS–NITHI methods. (related to Table S1)

| Source | SS | df | MS | F | Prob>F |
| --- | --- | --- | --- | --- | --- |
| Groups | $3.338 \times 10^4$ | 7 | $4.768 \times 10^3$ | $2.081 \times 10^3$ | 0 |
| Error | $4.583 \times 10^5$ | 199992 | 2.291 | | |
| Total | $4.916 \times 10^5$ | 199999 | 0 | | |

**Table S5.** Post-hoc multiple comparison test of number of stimuli of IFCN and MILLS–NITHI methods. (related to Table S1)

| Group A | Group B | Mean (Group A – Group B) | 95% CI (Group A – Group B) |  | p-value |
| --- | --- | --- | --- | --- | --- |
| 1 | 2 | 53.30 | 52.95 | 53.64 | 0 |
| 1 | 3 | 68.54 | 68.20 | 68.89 | 0 |
| 1 | 4 | 93.44 | 93.10 | 93.79 | 0 |
| 1 | 5 | 68.40 | 68.05 | 68.75 | 0 |
| 1 | 6 | 82.53 | 82.18 | 82.87 | 0 |
| 1 | 7 | 71.00 | 70.65 | 71.34 | 0 |
| 1 | 8 | 86.01 | 85.66 | 86.35 | 0 |
| 2 | 3 | 15.25 | 14.90 | 15.59 | 0 |
| 2 | 4 | 40.15 | 39.80 | 40.49 | 0 |
| 2 | 5 | 15.10 | 14.76 | 15.45 | 0 |
| 2 | 6 | 29.23 | 28.88 | 29.58 | 0 |
| 2 | 7 | 17.70 | 17.35 | 18.05 | 0 |
| 2 | 8 | 32.71 | 32.36 | 33.06 | 0 |
| 3 | 4 | 24.90 | 24.55 | 25.25 | 0 |
| 3 | 5 | −0.15 | −0.49 | 0.20 | **0.91** |
| 3 | 6 | 13.98 | 13.64 | 14.33 | 0 |
| 3 | 7 | 2.45 | 2.11 | 2.80 | 0 |
| 3 | 8 | 17.46 | 17.12 | 17.81 | 0 |
| 4 | 5 | −25.04 | −25.39 | −24.70 | 0 |
| 4 | 6 | −10.92 | −11.26 | −10.57 | 0 |
| 4 | 7 | −22.45 | −22.79 | −22.10 | 0 |
| 4 | 8 | −7.44 | −7.78 | −7.09 | 0 |
| 5 | 6 | 14.13 | 13.78 | 14.47 | 0 |
| 5 | 7 | 2.60 | 2.25 | 2.94 | 0 |
| 5 | 8 | 17.61 | 17.26 | 17.95 | 0 |
| 6 | 7 | −11.53 | −11.88 | −11.18 | 0 |
| 6 | 8 | 3.48 | 3.13 | 3.83 | 0 |
| 7 | 8 | 15.01 | 14.66 | 15.36 | 0 |

**Table S6.** Post-hoc multiple comparison test of relative errors of IFCN and MILLS–NITHI methods. (related to Table S1)

| Group A | Group B | Mean (Group A – Group B) | 95% CI (Group A – Group B) |  | p-value |
| --- | --- | --- | --- | --- | --- |
| 1 | 2 | −0.30 | −0.36 | −0.24 | 0 |
| 1 | 3 | −0.23 | −0.29 | −0.16 | 0 |
| 1 | 4 | −0.22 | −0.28 | −0.16 | 0 |
| 1 | 5 | 0.97 | 0.91 | 1.04 | 0 |
| 1 | 6 | 1.05 | 0.99 | 1.11 | 0 |
| 1 | 7 | 1.11 | 1.05 | 1.17 | 0 |
| 1 | 8 | 1.11 | 1.05 | 1.17 | 0 |
| 2 | 3 | 0.08 | 0.02 | 0.14 | $4 \times 10^{-3}$ |
| 2 | 4 | 0.08 | 0.02 | 0.15 | $10^{-3}$ |
| 2 | 5 | 1.28 | 1.22 | 1.34 | 0 |
| 2 | 6 | 1.35 | 1.29 | 1.41 | 0 |
| 2 | 7 | 1.41 | 1.35 | 1.48 | 0 |
| 2 | 8 | 1.41 | 1.35 | 1.48 | 0 |
| 3 | 4 | 0.01 | −0.06 | 0.07 | **1** |
| 3 | 5 | 1.20 | 1.14 | 1.26 | 0 |
| 3 | 6 | 1.27 | 1.21 | 1.33 | 0 |
| 3 | 7 | 1.34 | 1.28 | 1.40 | 0 |
| 3 | 8 | 1.34 | 1.28 | 1.40 | 0 |
| 4 | 5 | 1.19 | 1.13 | 1.26 | 0 |
| 4 | 6 | 1.27 | 1.20 | 1.33 | 0 |
| 4 | 7 | 1.33 | 1.27 | 1.39 | 0 |
| 4 | 8 | 1.33 | 1.27 | 1.39 | 0 |
| 5 | 6 | 0.07 | 0.01 | 0.13 | $9 \times 10^{-3}$ |
| 5 | 7 | 0.14 | 0.08 | 0.20 | $4 \times 10^{-10}$ |
| 5 | 8 | 0.14 | 0.08 | 0.20 | $4 \times 10^{-10}$ |
| 6 | 7 | 0.06 | 0.00 | 0.13 | 0.03 |
| 6 | 8 | 0.06 | 0.00 | 0.13 | 0.03 |
| 7 | 8 | 0.00 | −0.06 | 0.06 | 1 |

**Table S7.**
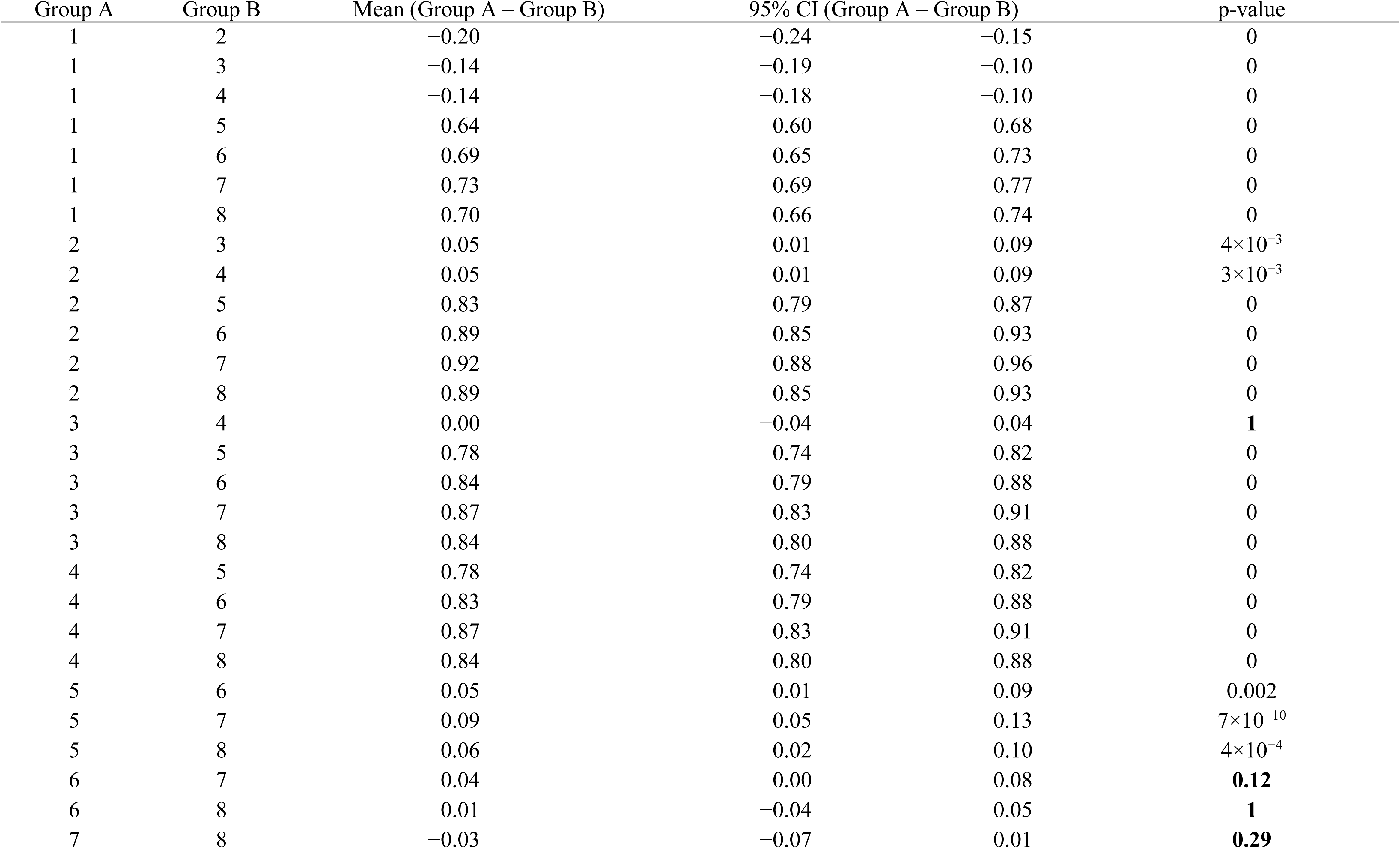
Post-hoc multiple comparison test of absolute errors of IFCN and MILLS–NITHI methods. (related to Table S1)

| Group A | Group B | Mean (Group A – Group B) | 95% CI (Group A – Group B) |  | p-value |
| --- | --- | --- | --- | --- | --- |
| 1 | 2 | −0.20 | −0.24 | −0.15 | 0 |
| 1 | 3 | −0.14 | −0.19 | −0.10 | 0 |
| 1 | 4 | −0.14 | −0.18 | −0.10 | 0 |
| 1 | 5 | 0.64 | 0.60 | 0.68 | 0 |
| 1 | 6 | 0.69 | 0.65 | 0.73 | 0 |
| 1 | 7 | 0.73 | 0.69 | 0.77 | 0 |
| 1 | 8 | 0.70 | 0.66 | 0.74 | 0 |
| 2 | 3 | 0.05 | 0.01 | 0.09 | $4 \times 10^{-3}$ |
| 2 | 4 | 0.05 | 0.01 | 0.09 | $3 \times 10^{-3}$ |
| 2 | 5 | 0.83 | 0.79 | 0.87 | 0 |
| 2 | 6 | 0.89 | 0.85 | 0.93 | 0 |
| 2 | 7 | 0.92 | 0.88 | 0.96 | 0 |
| 2 | 8 | 0.89 | 0.85 | 0.93 | 0 |
| 3 | 4 | 0.00 | −0.04 | 0.04 | **1** |
| 3 | 5 | 0.78 | 0.74 | 0.82 | 0 |
| 3 | 6 | 0.84 | 0.79 | 0.88 | 0 |
| 3 | 7 | 0.87 | 0.83 | 0.91 | 0 |
| 3 | 8 | 0.84 | 0.80 | 0.88 | 0 |
| 4 | 5 | 0.78 | 0.74 | 0.82 | 0 |
| 4 | 6 | 0.83 | 0.79 | 0.88 | 0 |
| 4 | 7 | 0.87 | 0.83 | 0.91 | 0 |
| 4 | 8 | 0.84 | 0.80 | 0.88 | 0 |
| 5 | 6 | 0.05 | 0.01 | 0.09 | 0.002 |
| 5 | 7 | 0.09 | 0.05 | 0.13 | $7 \times 10^{-10}$ |
| 5 | 8 | 0.06 | 0.02 | 0.10 | $4 \times 10^{-4}$ |
| 6 | 7 | 0.04 | 0.00 | 0.08 | **0.12** |
| 6 | 8 | 0.01 | −0.04 | 0.05 | **1** |
| 7 | 8 | −0.03 | −0.07 | 0.01 | **0.29** |

## Supplementary figures

**Figure S1:**
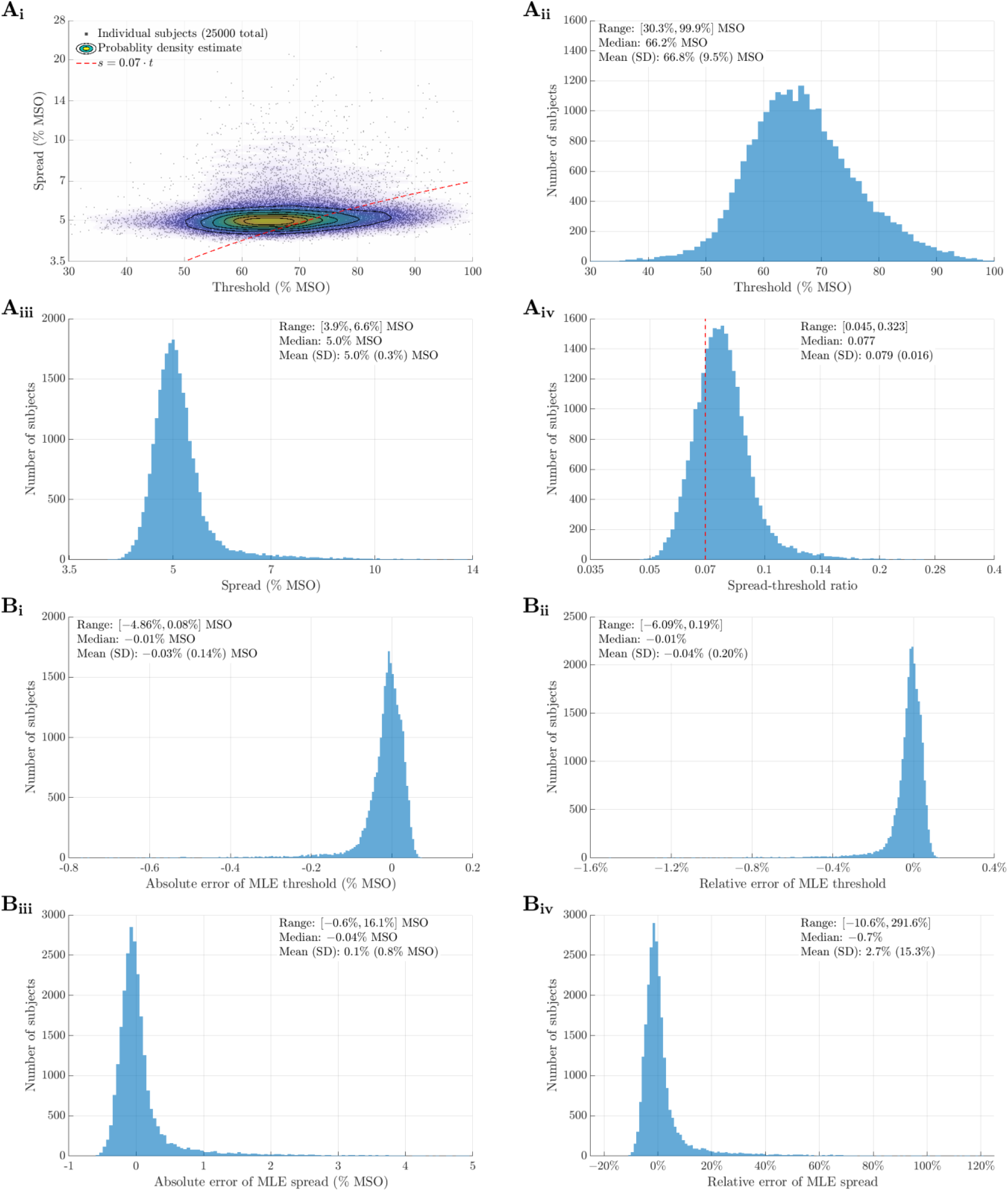
Maximum likelihood estimation (MLE) thresholds and slopes of model population. **A**. Distribution of MLE thresholds *t* and spread *s*, similar to Figure 1B. **B.** Distributions of absolute and relative errors of MLE threshold and spread compared to ground truth obtained from linear regression.

**Figure S2:**
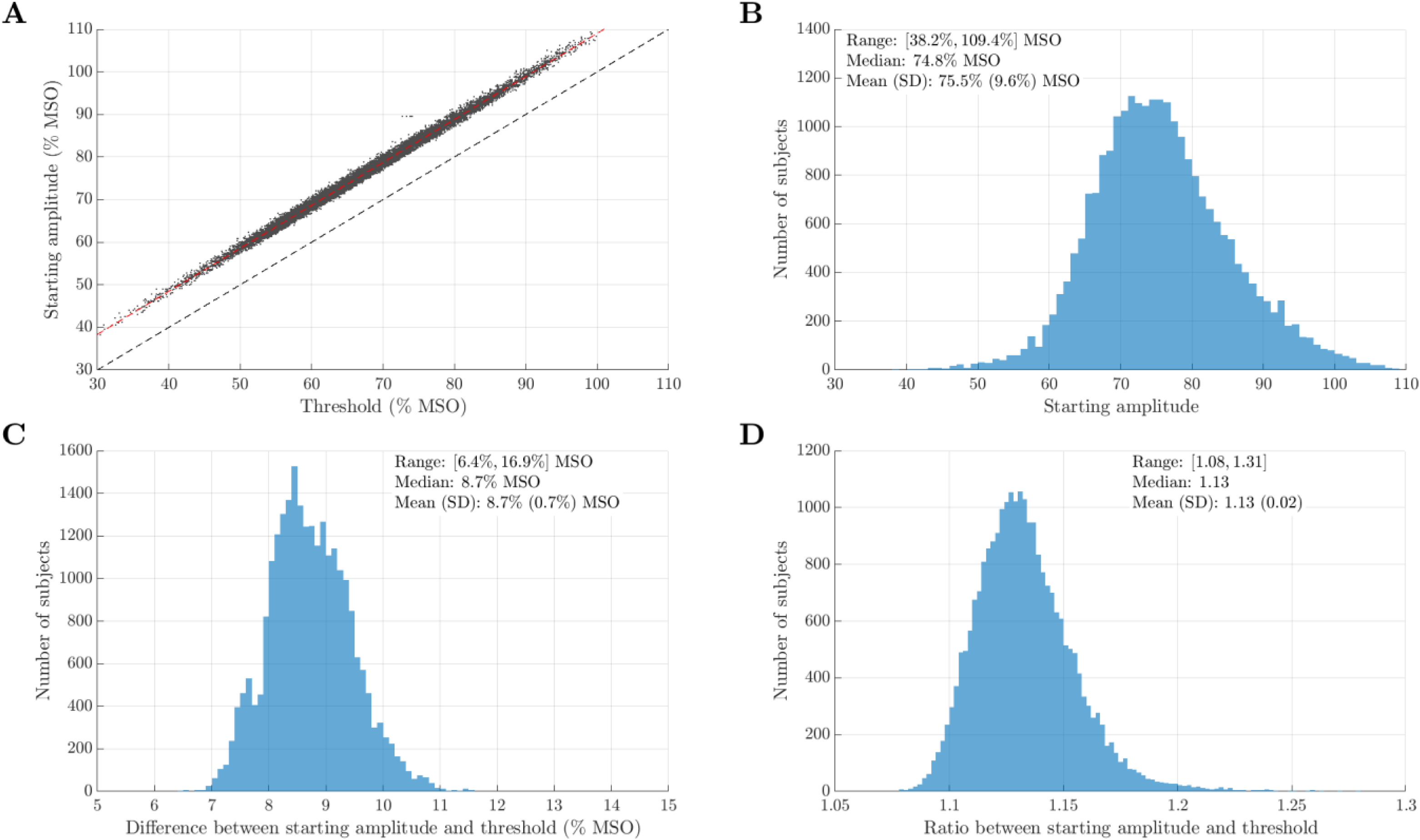
Starting amplitudes. **A**. Distribution of the starting amplitudes versus ground truth threshold, with a highly linear relationship (coefficient of determination *r*^2^>0.99). **B.** Distribution of the starting amplitudes. **C.** Difference of starting amplitudes and ground truth threshold, centered at 8.7% MSO. **D.** Ratio of starting amplitudes and ground truth threshold.

**Figure S3:**
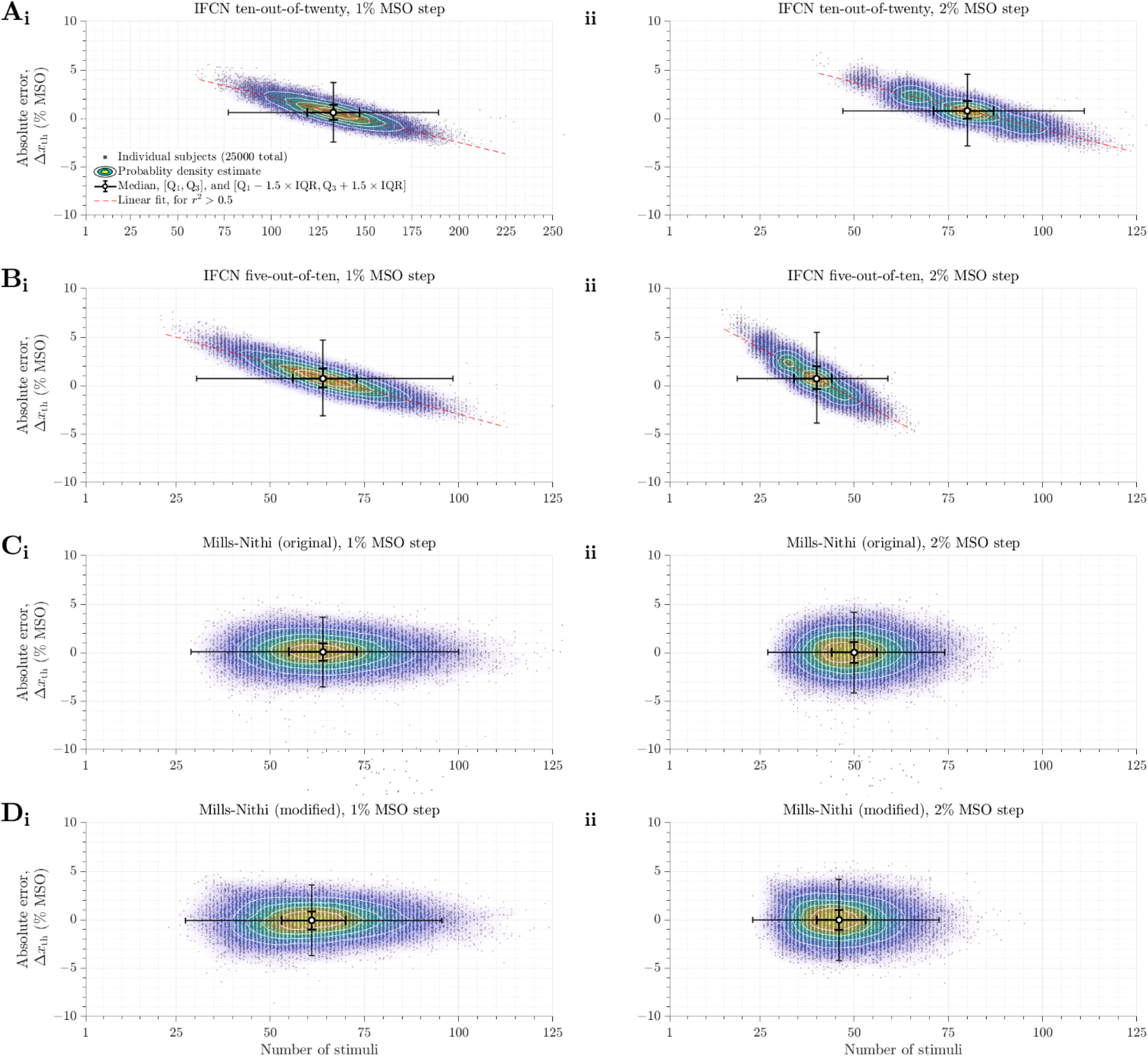
Absolute threshold errors of the IFNC relative frequency and MILLS–NITHI methods. Shown similarly as the relative threshold errors in Figure 2.

**Figure S4:**
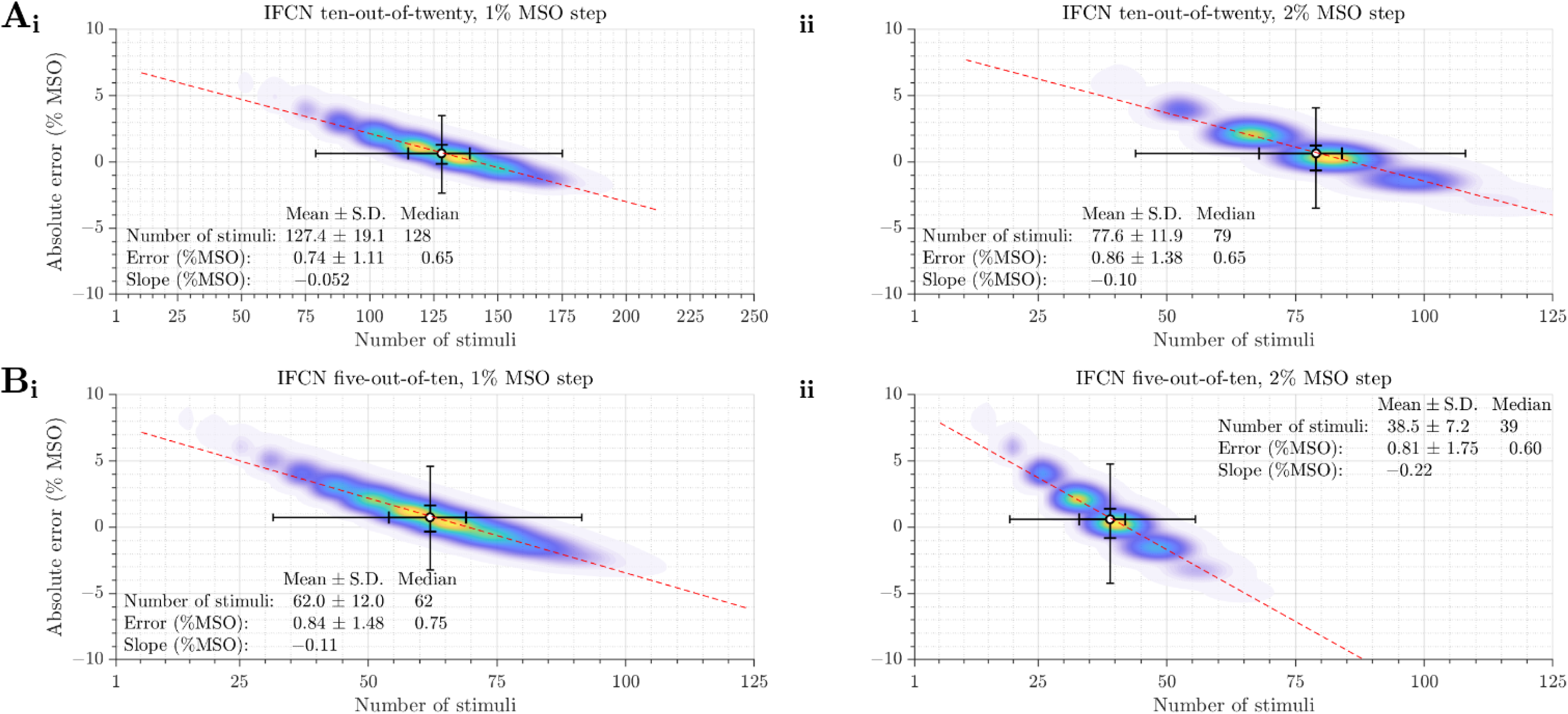
Theoretical distribution of absolute threshold errors of the IFNC relative-frequency methods. Similar to Figure S3.

**Figure S5:**
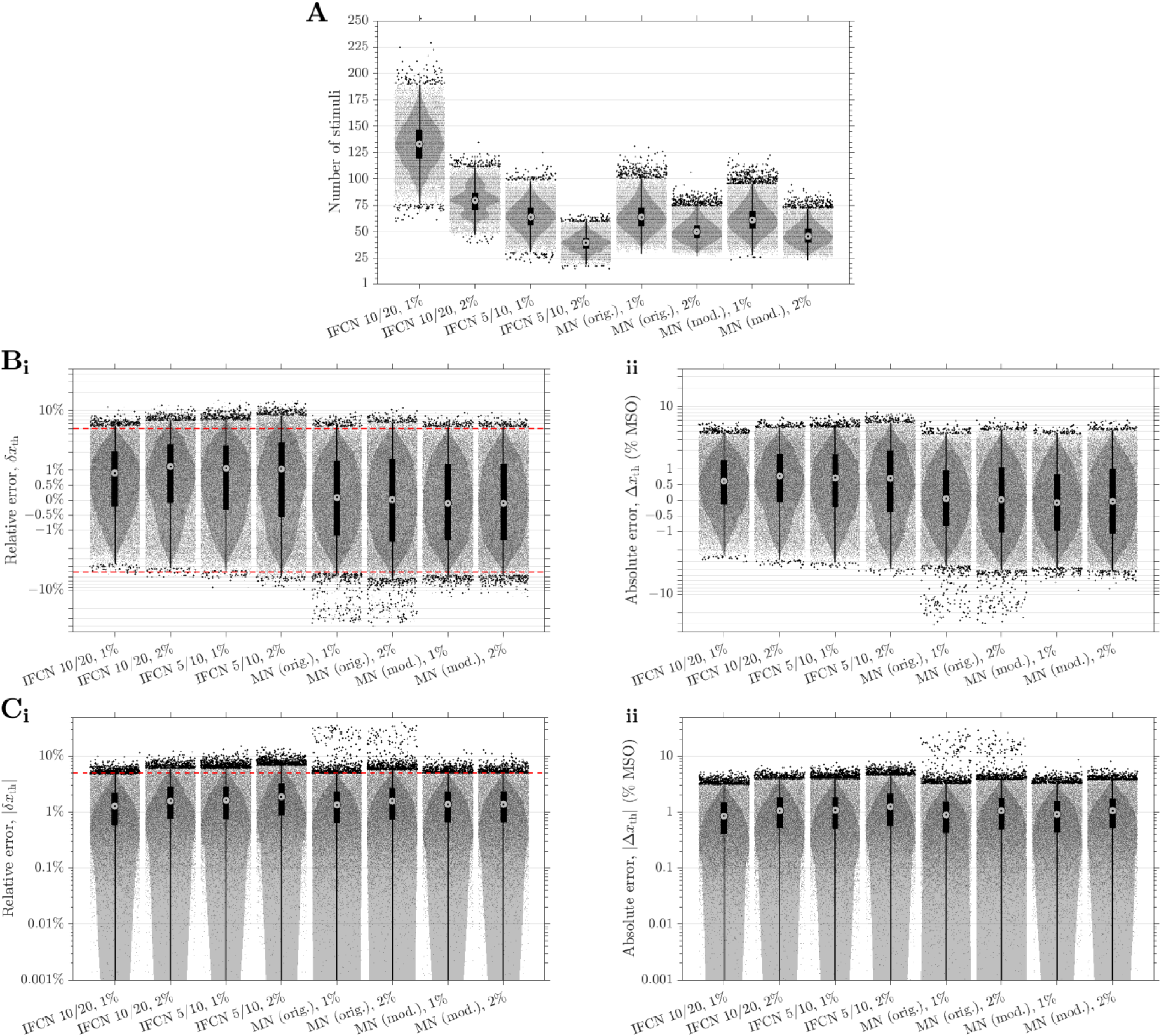
Statistics of IFCN relative frequency and MILLS–NITHI methods. The box-and-whisker plots show the medians (circles), interquartile ranges (thick lines). The ranges extending to the lower and upper adjacent values—1.5 times the interquartile range below and above the first and third quartiles (Q1 and Q3), respectively—are shown as whiskers (thin lines). All individual data points are shown, with outliers as darker dots. The gray patches are probability density estimates based on a normal kernel function. **A.** Number of stimuli. **B.** Threshold errors shown on a log-lin-log mixed scale to better visualize the errors around zero, with linear scaling in the range of [−1%, 1%] and [−1, 1] %MSO for the relative and absolute errors, respectively, and logarithmic scaling of amplitudes outside these ranges. **i.** Relative error of the threshold. **ii.** Absolute error of the threshold. **C.** Absolute value of threshold errors shown on logarithmic scale. The whiskers and probability density estimates extend to zero, but are cut off at 0.001% and 0.001% MSO, respectively, for visualization. **i.** Relative error of threshold. **ii.** Absolute error of threshold. The ±5% criteria for safety are shown as red dashed line for the relative errors in panels Bi and Ci.

**Figure S6:**
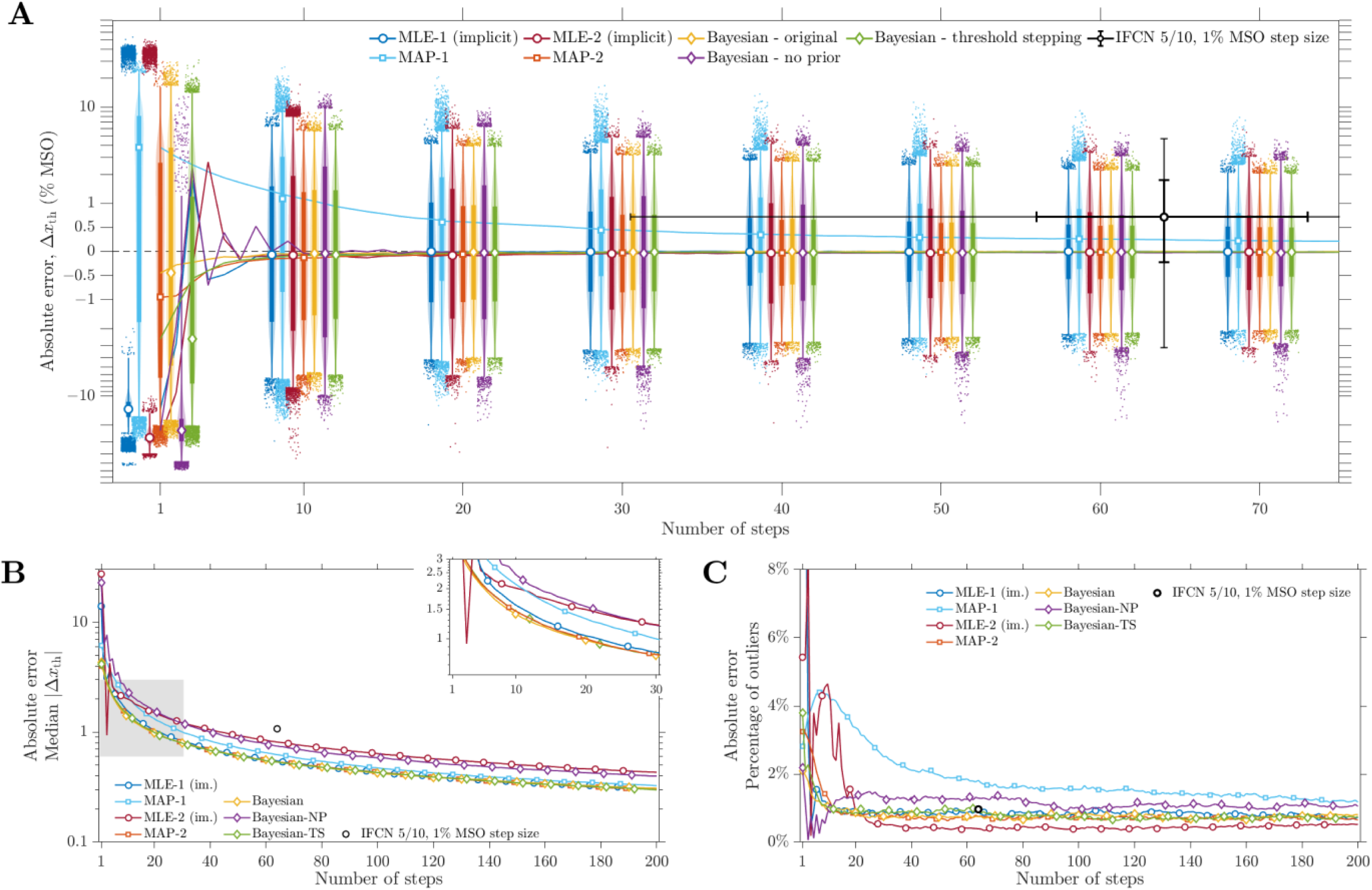
Absolute threshold errors of parametric estimation methods. Shown similarly as the relative threshold error in Figure 3.

**Figure S7:**
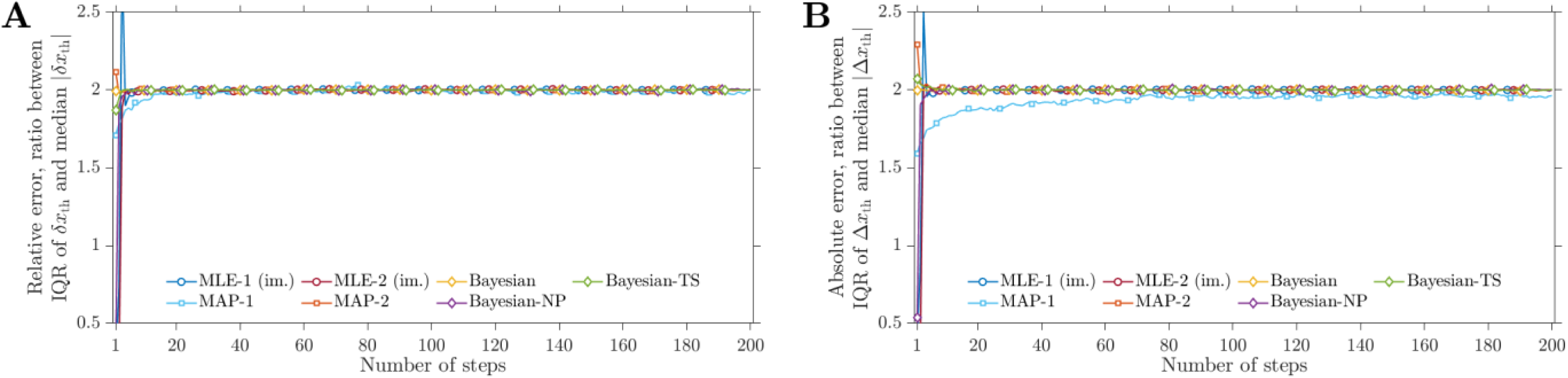
Ratio of median of absolute value of errors and interquartile ranges (IQR) of errors of parametric estimation methods. **A**. Ratio for relative threshold errors (related to Figures 3). **B.** Ratio for absolute threshold errors (related to Figure S6).

**Figure S8:**
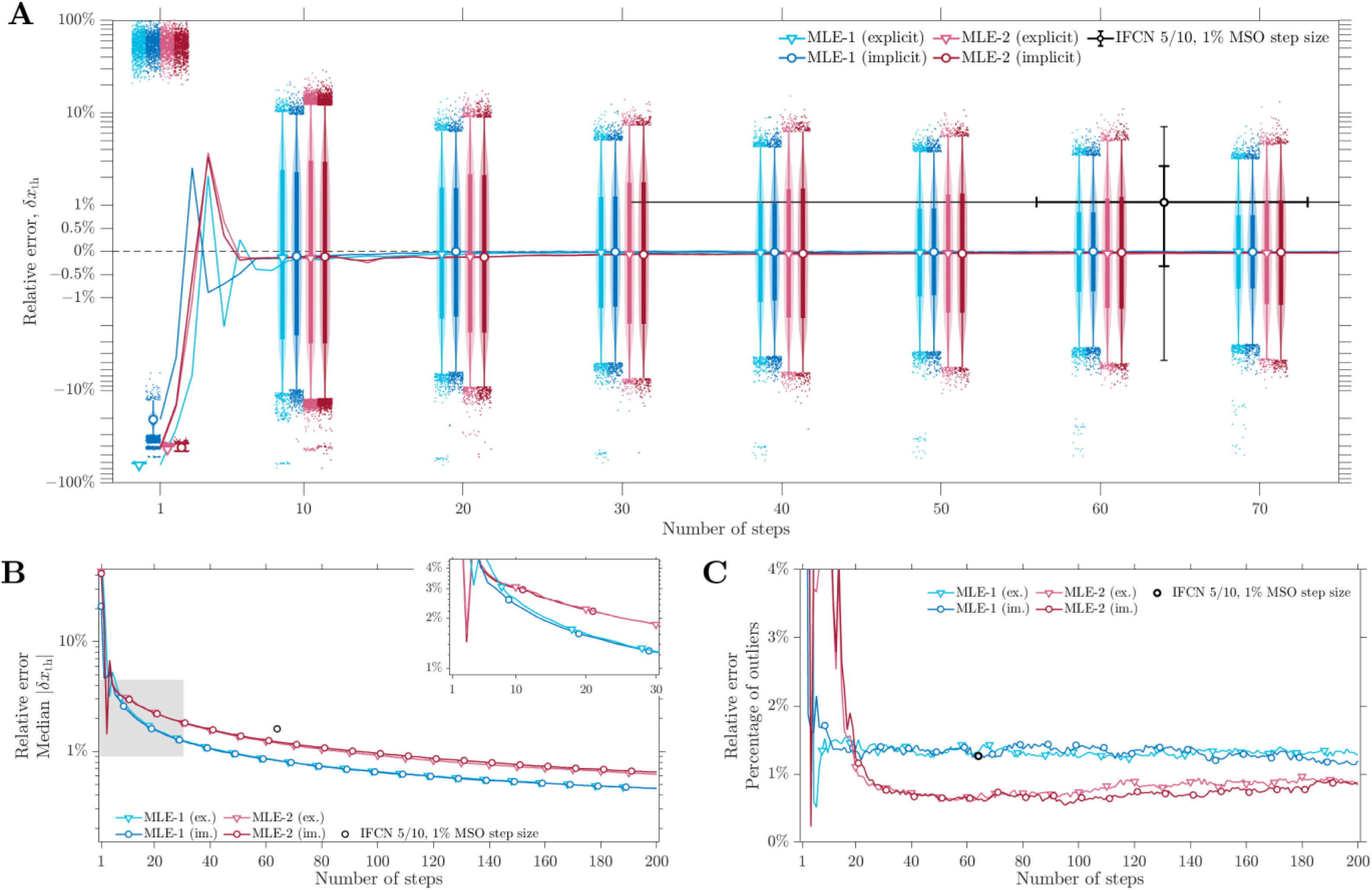
Relative threshold errors of MLE methods using explicit (ex.) and implicit (im.) maximization of the likelihood function. Shown similarly as in Figure 3.

**Figure S9:**
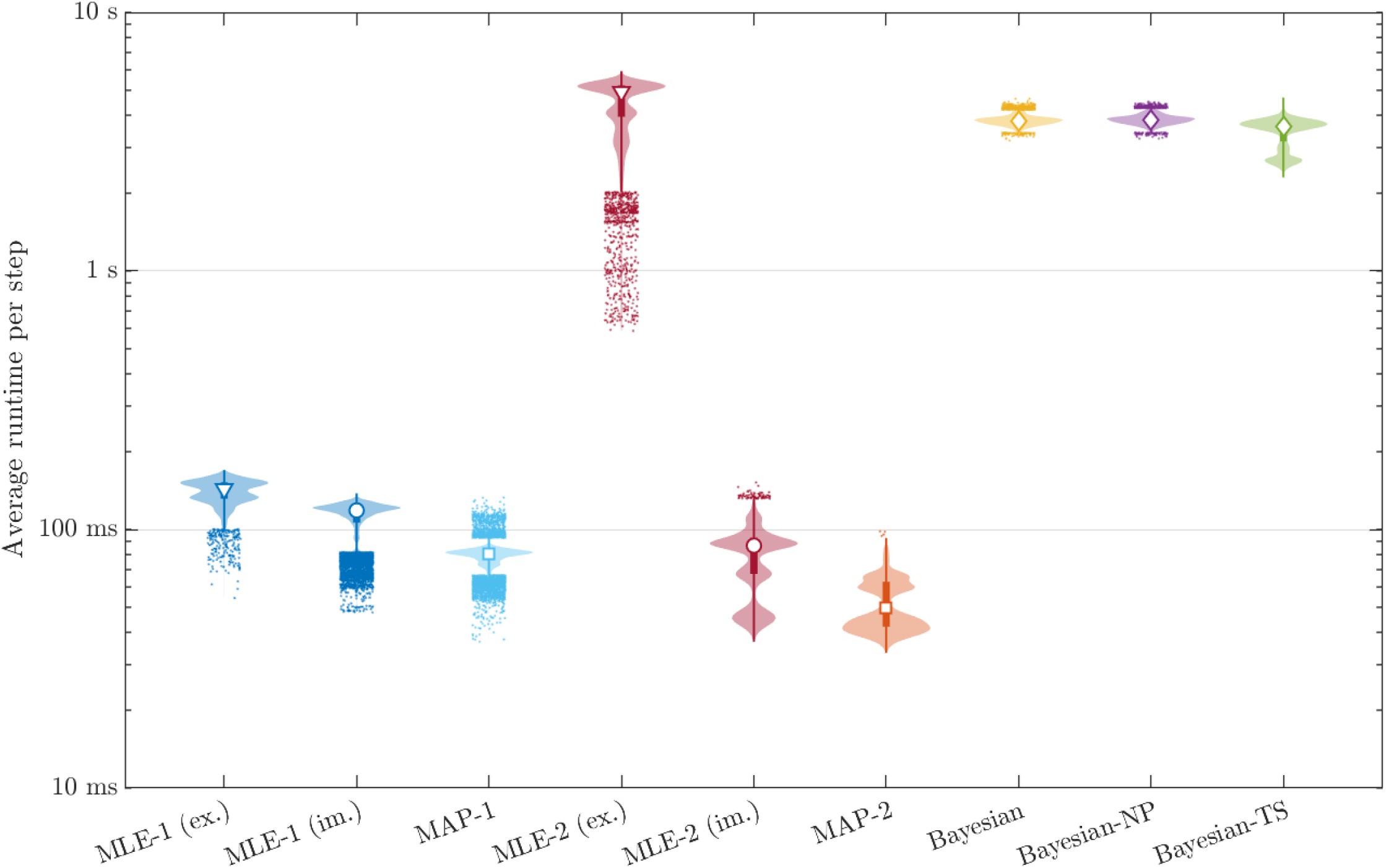
Runtime of parametric estimation methods. The distribution of average runtime per step shown for the 25,000 virtual subjects.

**Figure S10:**
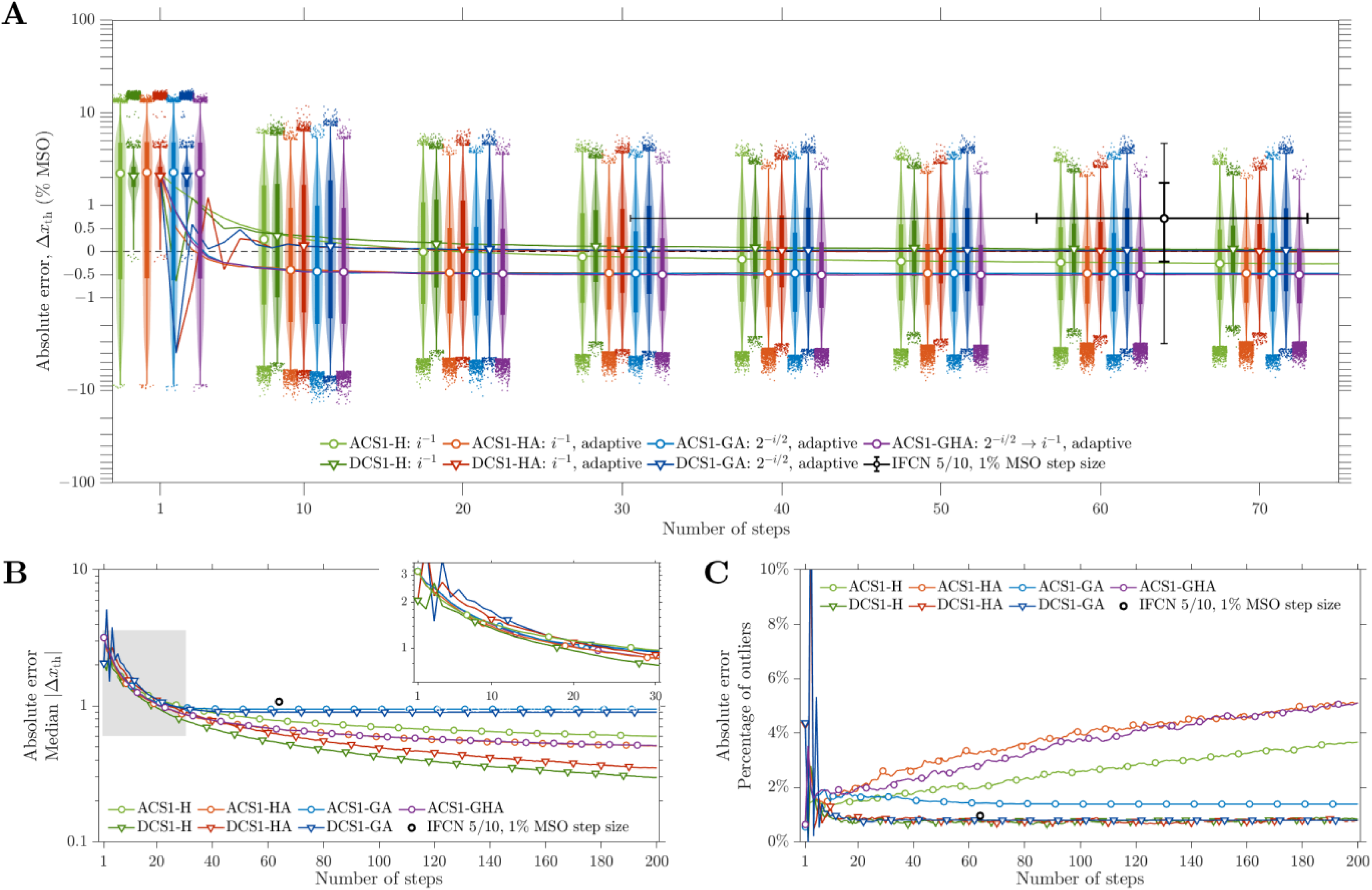
Absolute threshold errors of root finding methods using first-order control sequence with default initial step size a_0_. Shown similarly as the relative threshold error in Figure 4.

**Figure S11:**
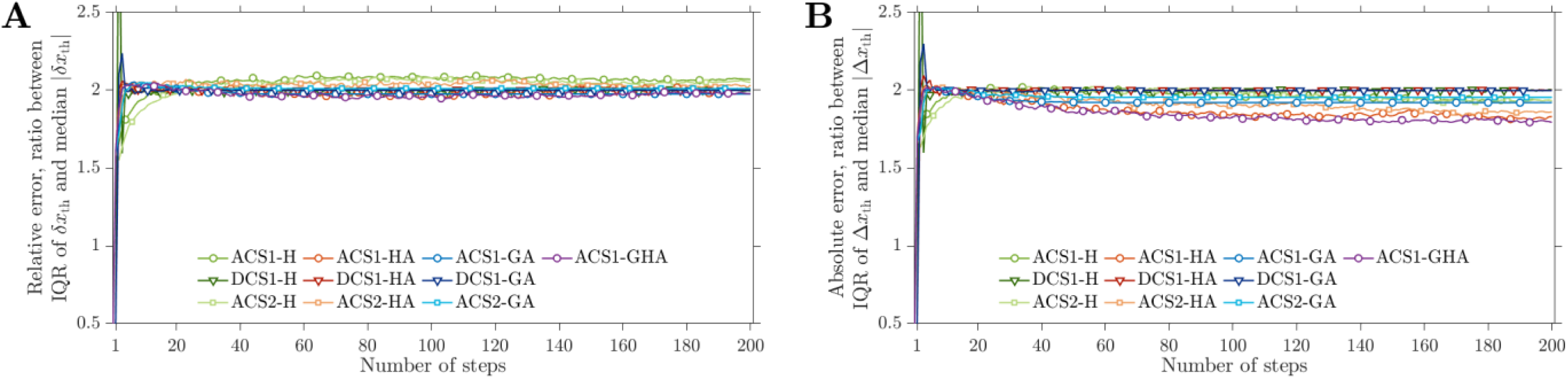
Ratio of median of absolute value of errors and interquartile range (IQR) of errors of root finding methods using default parameters. **A.** Ratio for relative threshold errors (related to Figures 4 and S15A). **B.** Ratio for absolute threshold errors (related to Figures S10 and S15B).

**Figure S12:**
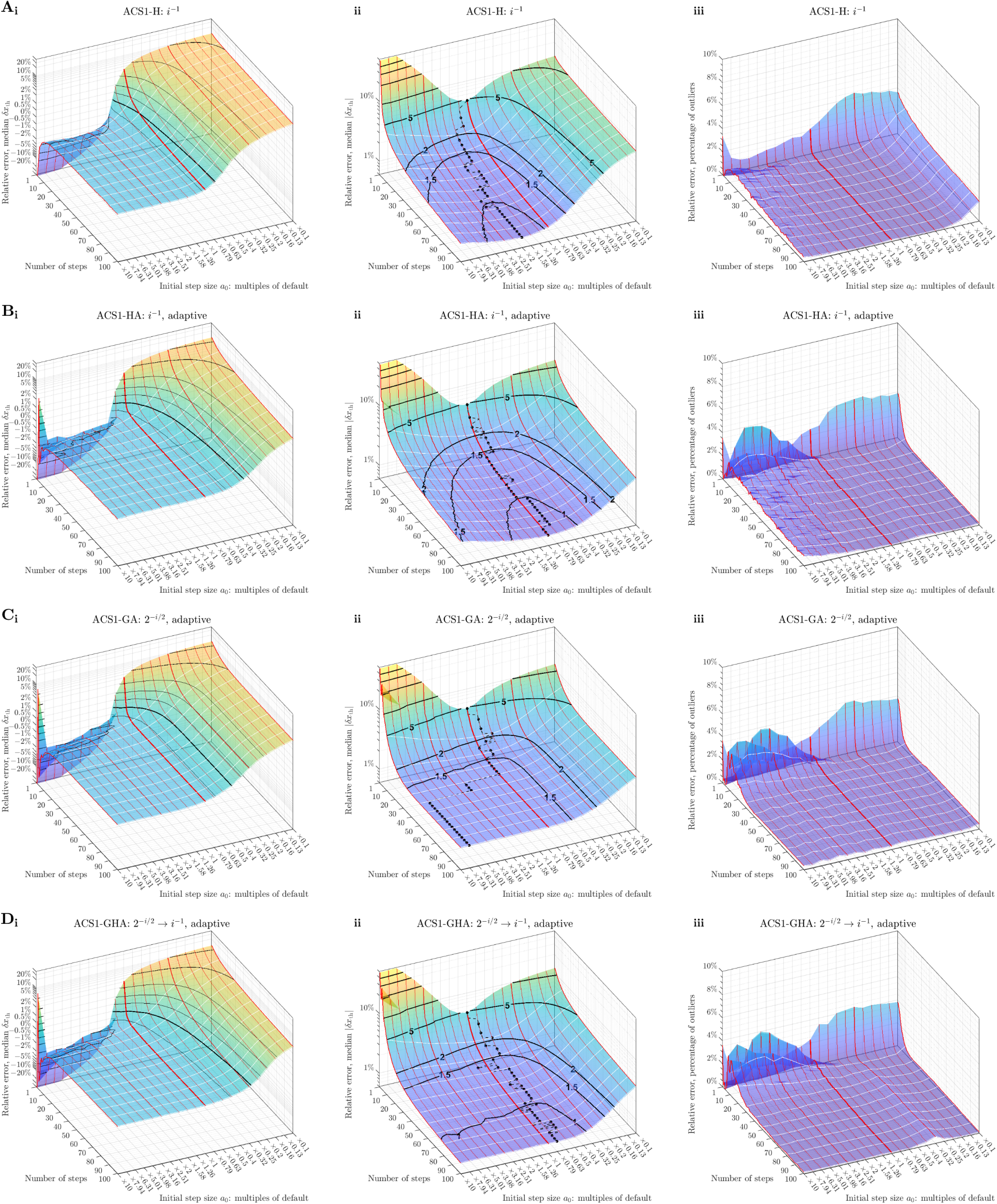
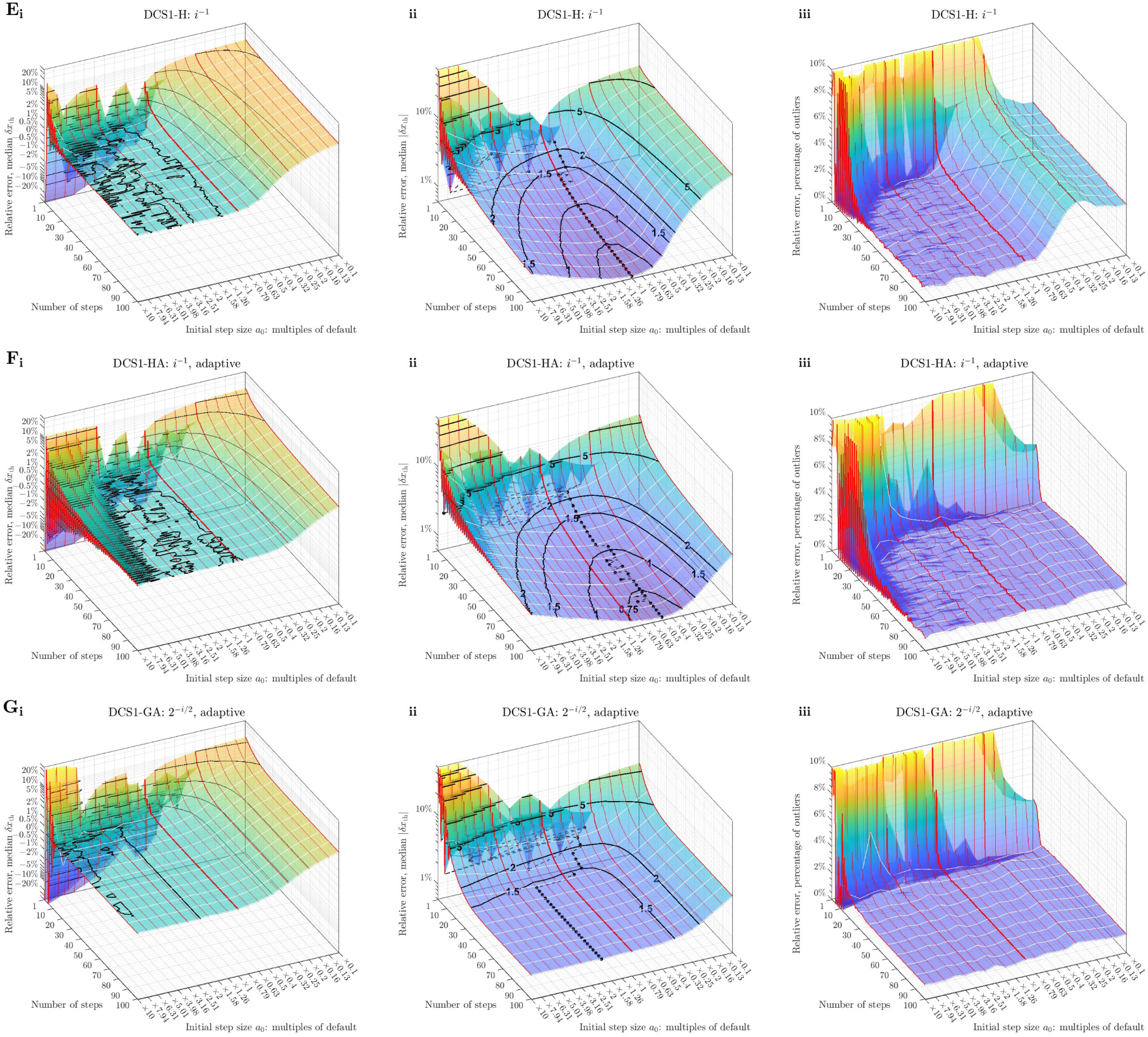
Relative threshold errors of root finding methods using first-order control sequence for different initial step size a_0_. The red lines show results for different initial step size *a*_0_, which are provided as multiples of the default value, with the thick red line showing the default value (results in Figures 4). The white lines mark every tenth step. **A.** Results for ACS1-H. **i.** Median of errors shown on log-lin-log scale (as in Figures 4A). Black contour lines are shown at levels corresponding to the numbered ticks on the *z*-axis, with the thick line indicating zero. **ii.** Median of absolute value of errors shown on logarithmic scale (as in Figures 4B). The minimum at every step is shown with the black dashed line with a black dot at every third step. **iii.** Percentage of outliers (as in Figures 4C). **B.**–**D.** Results for ACS1-HA, ACS1-GA, and ACS1-GHA, respectively. **E.**–**G.** Results for DCS1-H, DACS1-HA, and DCS1-GA, respectively.

**Figure S13:**
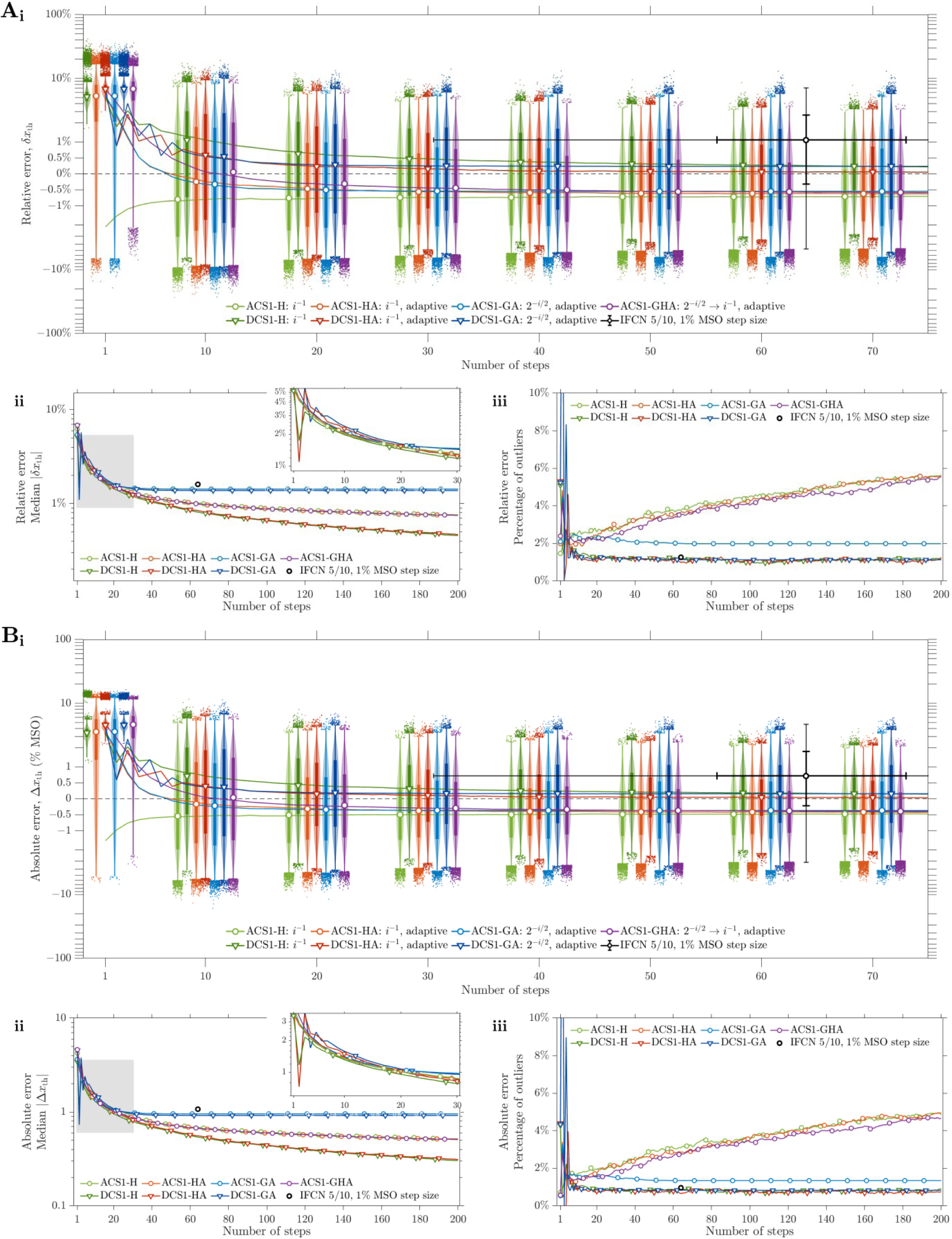
Threshold errors of root finding methods using first-order control sequence with optimal initial step size a_0_. Shown similarly as in Figures 4 and S10. **A.** Relative threshold error. **B.** Absolute threshold error.

**Figure S14:**
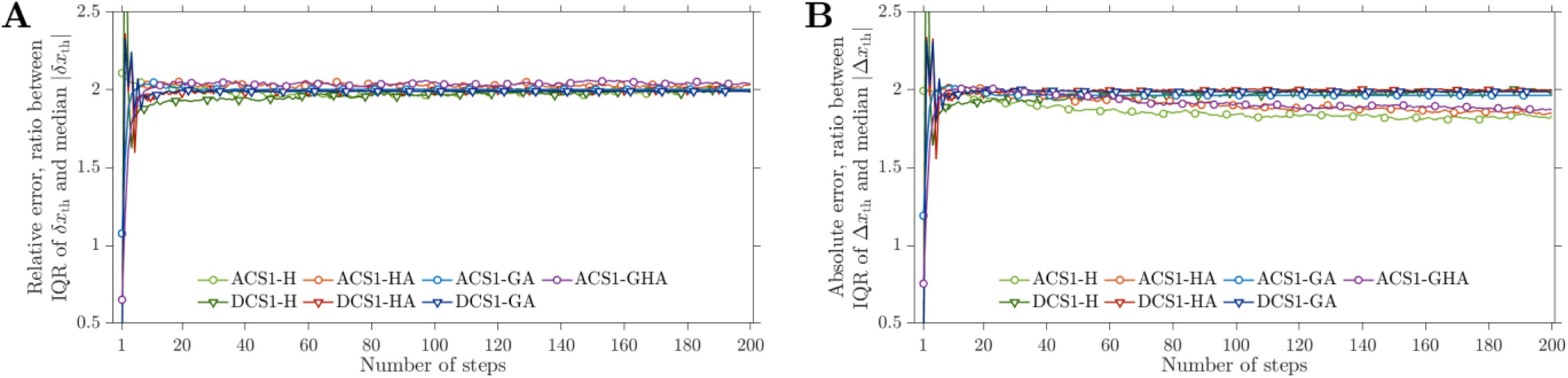
Ratio of median of absolute value of errors and interquartile range (IQR) of errors of first-order root finding methods using optimal initial step size a_0_. **A.** Ratio for relative threshold errors (related to Figures S13A). **B.** Ratio for absolute threshold errors (related to Figures S13B).

**Figure S15:**
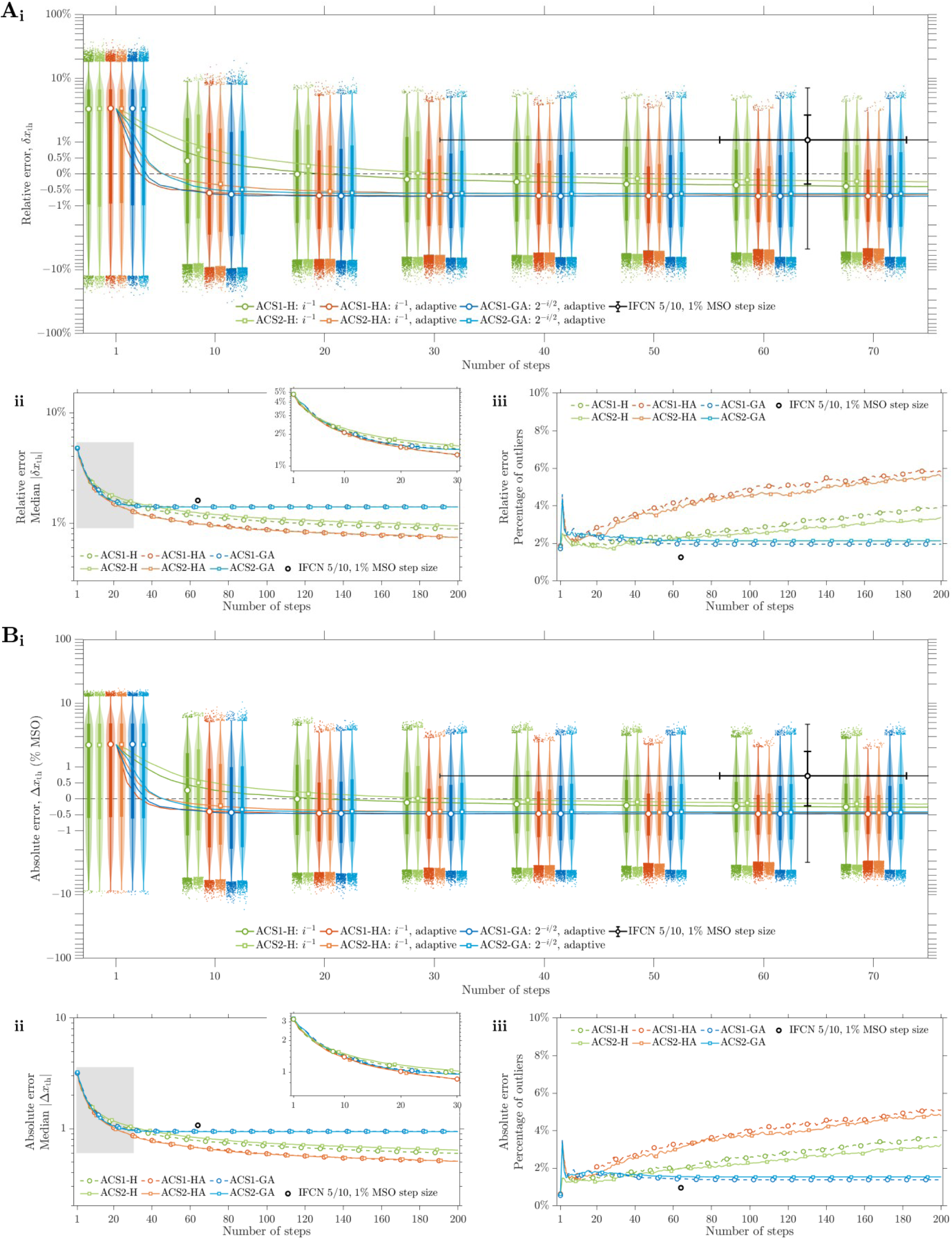
Threshold errors of root finding methods using second-order analog control sequence with default parameters. The corresponding methods using first-order analog control sequence are also shown for comparison (as in Figures 4 and S10). **A.** Relative threshold error. **B.** Absolute threshold error.

**Figure S16:**
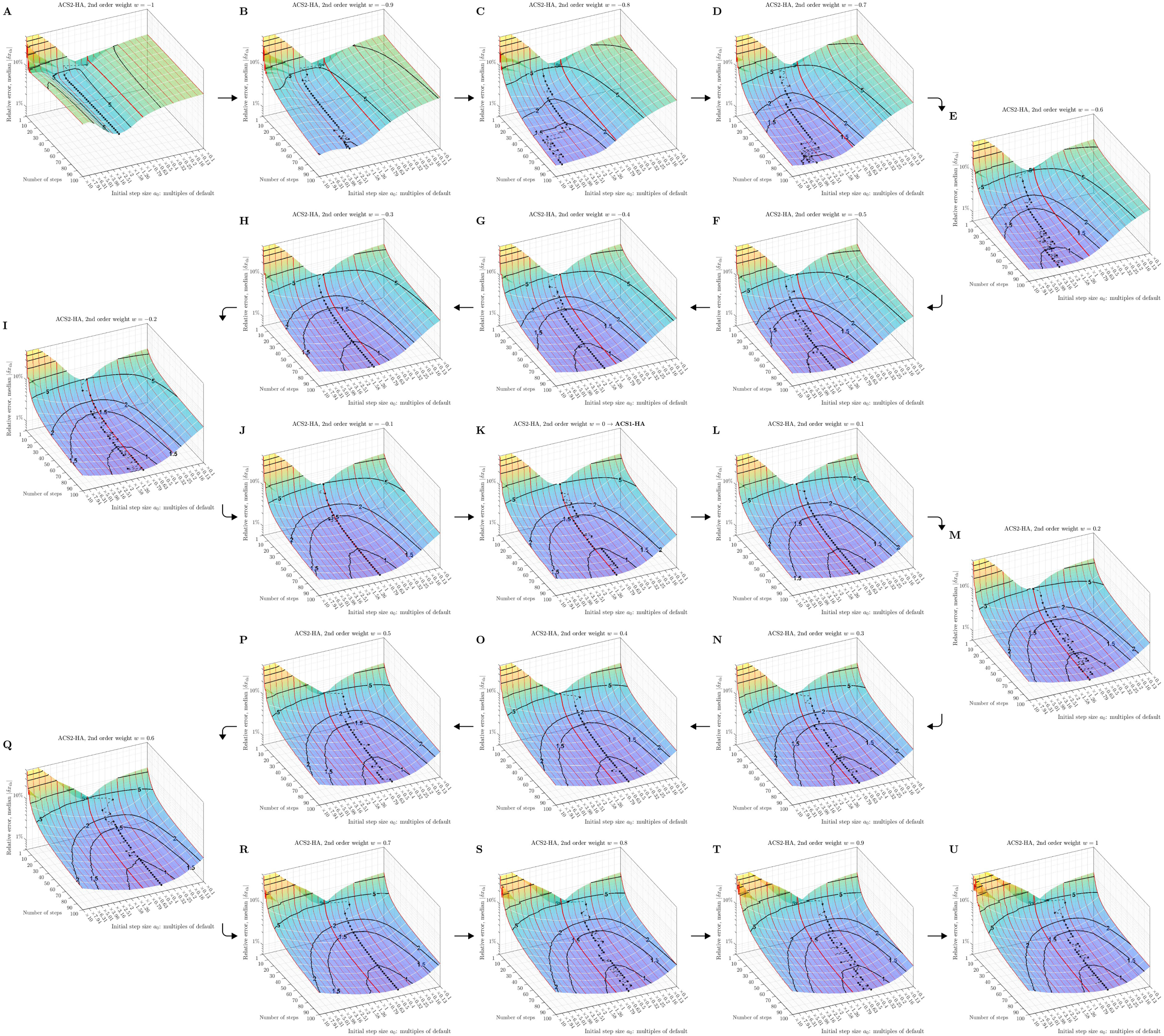
Relative threshold errors of root finding methods using second-order analog control sequence for different initial step size a_0_ and given second order weights *w*. **A.**–**U.** Results for second order weights from −1 to 1 in steps of 0.1, shown similarly as in Figure S12Aii.

**Figure S17:**
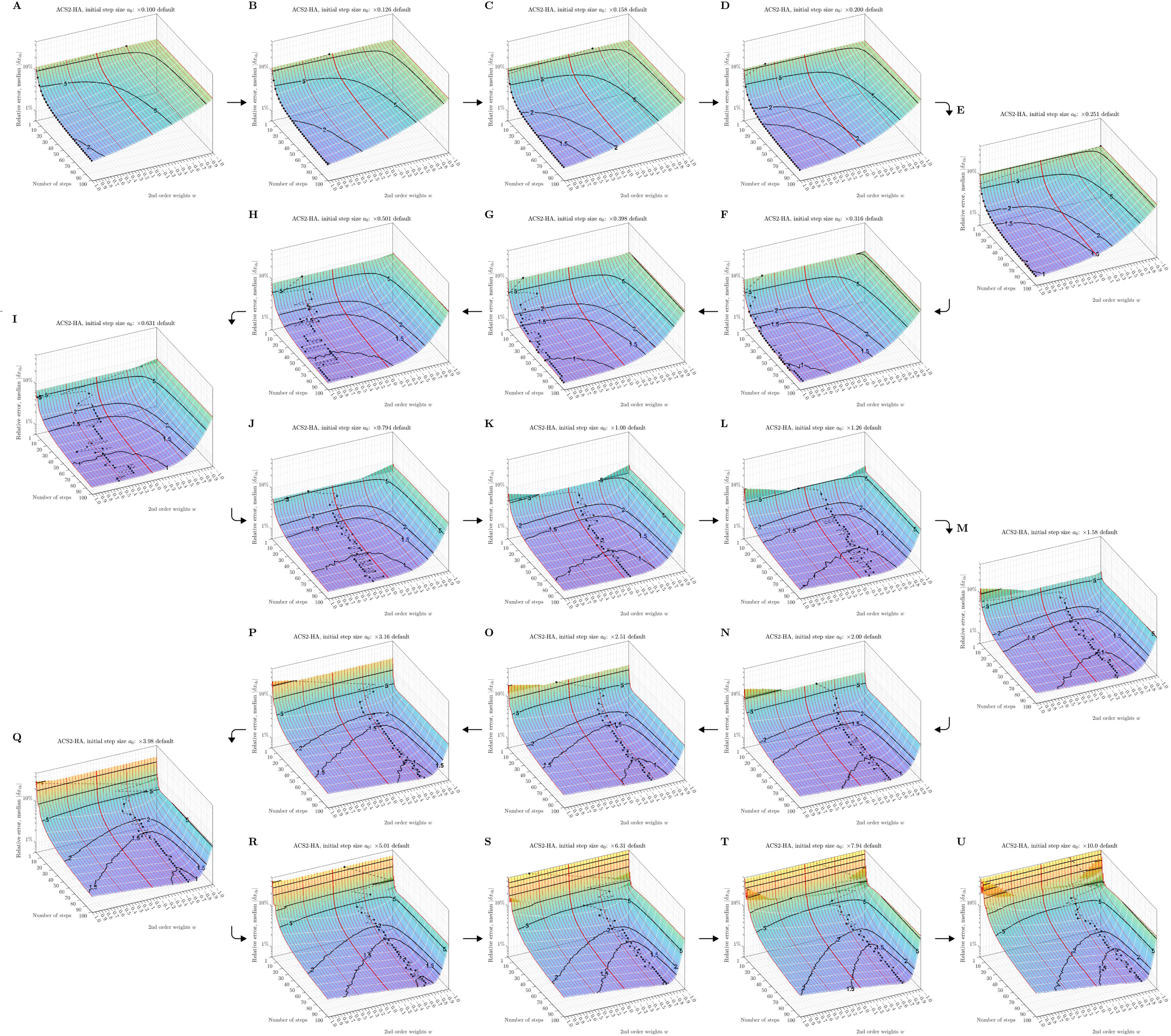
Relative threshold errors of root finding methods using second-order analog control sequence for different second order weights *w* and given initial step size a_0_. **A.**–**U.** Results for initial step size from 0.1 times to 10 times the default value, uniformly sampled on logarithmic scale, shown similarly as Figure S12Aii but with second-order weight *w* as the parameter. The thick red line indicates *w*=0, corresponding to the first-order version.

**Figure S18:**
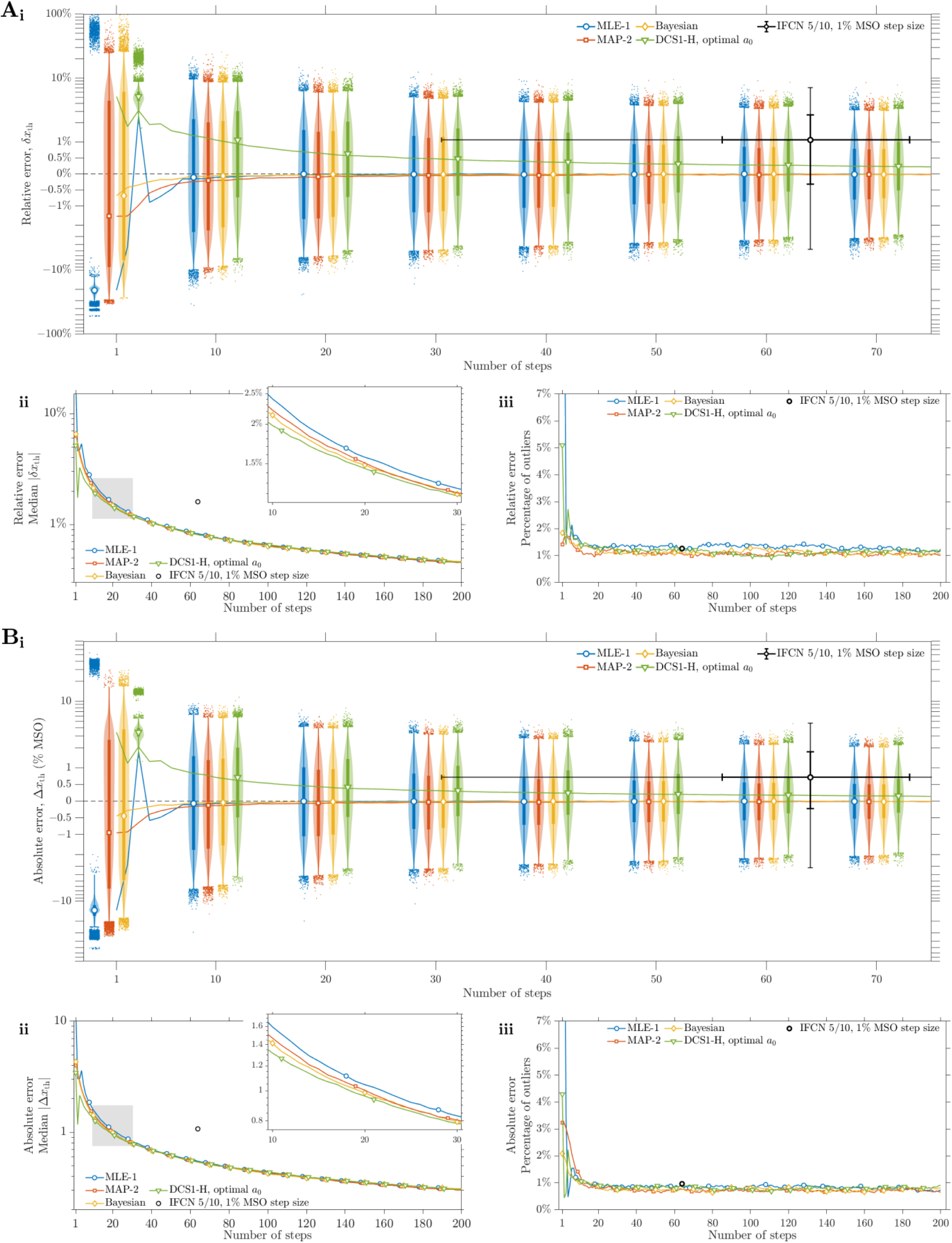
Threshold errors of the best performing methods in each category Related to Figure 5. Similar format as Figures 3 and S6. **A.** Relative threshold error. **B.** Absolute threshold error.

## Footnotes

1 The midpoint or threshold determines the horizontal location of the distribution and the slope at threshold or spread determines its shape, i.e., horizontal stretching. When referring to the distribution’s shape parameter, slope and spread are sometimes used interchangeably, although they are inversely related to each other.

1 Defined as convergence with probability one: 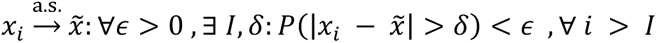

## Notes

https://doi.org/10.5281/zenodo.6483601

### Summary of Updates

Improved methods and updated results.

